# Over-representation of sperm-associated deleterious mutations across wild and *ex situ* cheetah (*Acinonyx jubatus*) populations

**DOI:** 10.64898/2026.04.07.716683

**Authors:** Jessica A. Peers, Heather R. Sibley, Ellie E. Armstrong, Adrienne E. Crosier, Will J. Nash, Klaus-Peter Koepfli, Wilfried Haerty

**Affiliations:** Earlham Institute, Norwich Research Park, Norwich, UK; College of Science, George Mason University, Fairfax, VA, USA; University of California, Riverside, Riverside, CA, USA; Center for Animal Care Sciences, Smithsonian’s National Zoo & Conservation Biology Institute, Front Royal, VA, USA; School of Biological Sciences, University of East Anglia, Norwich, UK; Colossal Biosciences, Dallas, TX, USA

**Keywords:** population genetics, *ex situ*, cheetah, inbreeding, effective population size, male fertility

## Abstract

As purifying selection becomes less effective and inbreeding increases, small populations frequently develop an increased load of genome-wide deleterious mutations. Reflecting this pattern, deleterious mutations in genes associated with fertility and immunity have previously been identified in the cheetah (*Acinonyx jubatus*), which has had a low effective population size for at least the last 10,000 years. However, the distribution of deleterious mutations across cheetah populations is currently unknown. Here, we analysed novel whole genome resequencing data from 30 *ex situ* and 9 wild cheetahs. We investigated variation in genetic diversity, genomic measures of inbreeding, and the distribution of deleterious mutations across cheetah populations. South Sudanese and Tanzanian cheetahs showed higher inbreeding and realized load, while Namibian cheetahs had a higher proportion of population-specific deleterious mutations. Genes containing high- or moderate-impact deleterious mutations were significantly enriched for sperm-related functions, highlighting putative causative loci associated with poor sperm quality in cheetahs. Similar levels of genetic diversity and inbreeding were observed in *ex situ* cheetahs compared to their wild counterparts, providing empirical evidence of the efficacy of captive breeding programmes in maintaining genetic variation in *ex situ* populations.

## Introduction

Small populations are prone to experiencing increased inbreeding and genetic drift resulting in an accumulation of deleterious mutations (Björnerfeldt et al., 2006; Lande, 1994; Masel, 2011). In wild populations, this can lead to ’extinction vortex’ effects, where fixation of deleterious variants and increases in inbreeding continually interact in a feedback loop, resulting in population extinction (Gilpin & Soulé, 1986). However, *ex situ* populations are intensively managed to ensure stable population sizes whilst minimising inbreeding. Despite the potential for such management to mitigate inbreeding effects, *ex situ* populations often originate from few individuals, resulting in strong founder effects, which decrease genetic diversity and increase inbreeding (Björnerfeldt et al., 2006; Muller, 1964). Additionally, in *ex situ* conditions, natural selection may be relaxed, allowing deleterious alleles to spread through the population (Lynch & O’Hely, 2001).

The cheetah (*Acinonyx jubatus*) presents an excellent opportunity to study the impacts of *ex situ* management on a species that has experienced significant inbreeding and prolonged low effective population size (*N_e_*). The cheetah is hypothesised to have experienced severe demographic decline beginning 10,000 years ago (Dobrynin et al., 2015; Fabiano et al., 2025; O’Brien & Johnson, 2007), resulting in sustained low *N_e_* and phenotypes linked with inbreeding depression, including morphological and skeletal defects, weakened immunity and decreased fertility (Castro-Prieto et al., 2011; O’Brien et al., 1985; Terio et al., 2018; Wayne et al., 1986; Wildt et al., 1983). Currently classified as ‘Vulnerable’ by the IUCN Red List of Threatened Species, approximately 7,000 cheetahs remain in the wild (Durant et al., 2017), with an estimated 2,000 cheetahs held in accredited facilities, excluding privately-owned animals (L. Marker & Johnston, 2022).

The first recorded cheetah in a modern zoological collection appeared in 1829 at the Zoological Society of London, though the first documented *ex situ* birth occurred in 1956 (Marker-Kraus, 1997). Although *ex situ* breeding efforts increased substantially starting in the 1970s, including the establishment of a coordinated programme by the Association of Zoos and Aquariums (AZA) in the 1980s, persistent reproductive challenges have limited their success (Marker-Kraus, 1997). Despite an *ex situ* population approaching 200 individuals, the effective breeding population was estimated at only 28, with infant mortality rates exceeding those observed in other species in *ex situ* settings (L. Marker & O’Brien, 1989). These reproductive limitations prompted a series of studies revealing low genetic diversity in the cheetah, particularly in MHC genes, as well as poor sperm quality (O’Brien et al., 1983, 1985; Wayne et al., 1986; Wildt et al., 1983).

By the late 1990s, the vast majority of known founders in the global *ex situ* population originated from the southern African subspecies (*A. j. jubatus*), with minimal representation from eastern Africa (*A. j. jubatus*, previously *A. j. raineyi*) and none from northern African (*A. j. soemmeringii* and *A. j. hecki*) or Asian (*A. j. venaticus*) subspecies (Marker-Kraus, 1997). Over 90% of *ex situ* cheetahs are believed to descend from wild-caught Namibian individuals, and although approximately 424 founders were part of the global *ex situ* population, 80% of those alive in 1994 had not yet bred successfully (Marker-Kraus, 1997).

While founder effects in the *ex situ* cheetah population may have been mitigated by consistent introduction of wild animals throughout the 20th century, such imports do not guarantee the maintenance of genetic diversity in the *ex situ* population. *Ex situ* breeding programmes select breeding pairs based on a minimization of kinship approach, typically informed by a pedigree (Couvet & Ronfort, 1994; Fernández & Caballero, 2001). However, whilst this practice may reduce breeding between close relatives, it does not necessarily reduce the spread and fixation of deleterious alleles that may already be present in the population due to the generally low *N_e_* of *ex situ* populations. Low *N_e_* can intensify genetic drift and decrease the efficiency of natural selection, increasing the risk of masked load (heterozygous deleterious mutations) converting to realized load (homozygous deleterious mutations) (Bertorelle et al., 2022; Lewin & Eyre-Walker, 2026; Smeds & Ellegren, 2023). Additionally, this method relies on the accuracy of the studbook, which is not well-maintained in all species (Giontella et al., 2020; Ivy & Lacy, 2010). Together, these factors can facilitate the accumulation and spread of deleterious variants within *ex situ* populations (Woodworth et al., 2002).

Cheetahs are known to exhibit signs of poor sperm quality and reproductive defects (O’Brien et al., 1985; Wildt et al., 1983), which may negatively impact reproduction in the wild. However, in *ex situ* conditions, individuals carrying such mutations may still reproduce, allowing the mutations to persist in the population. For example, *in vitro* fertilisation (IVF) was carried out on cheetahs in 2020, resulting in two offspring (Crosier et al., 2020). Whilst this use of IVF represents a significant advancement in the application of assisted-reproductive technologies for cheetah conservation, deleterious mutations which may have otherwise prohibited successful reproduction could be passed to offspring and maintained in the population.

The severe genetic bottleneck and low effective population size of cheetahs predates captivity (Barnett et al., 2005; Kim et al., 2016; O’Brien & Johnson, 2007; O’Regan & Steininger, 2017), meaning the majority of accumulated deleterious mutations are likely to be shared between wild and *ex situ* populations. However, these populations have been exposed to contrasting selection pressures and breeding histories. As *ex situ* breeding programmes often aim to act as insurance or reserve populations for potential rewilding and restoration efforts (Fraser, 2008; Witzenberger & Hochkirch, 2011), it is important to understand the relationship between the genetic diversity of wild populations and their *ex situ* equivalents.

Since the first observation of low genetic diversity in *ex situ* cheetahs (O’Brien et al., 1985), researchers have continued to investigate this phenomenon (Terrell et al., 2016). To date, the majority of studies have relied on microsatellite panels or targeted sequencing approaches such as double-digest restriction-site associated DNA sequencing (ddRAD-seq) (Castro-Prieto et al., 2011; Driscoll et al., 2002; Meißner et al., 2024; Prost et al., 2022; Terrell et al., 2016). While (Prost et al., 2022) generated the first genome-wide insights into genetic diversity in wild cheetah populations using a reduced representation approach, the limited availability of whole-genome data (n = 8) means that genome-wide patterns of diversity across populations remain largely unresolved (Dobrynin et al., 2015; Winter et al., 2023). In particular, genome-wide genetic diversity and load have not yet been characterised for any *ex situ* cheetah population.

Here, we sequenced and analysed whole genomes of 30 cheetahs from the *ex situ* population in the United States (U.S.) and two wild cheetahs, integrating these with existing genomic data from wild populations to investigate genetic structure, diversity, and the distribution of masked and realized deleterious variants. A consistent separation of *ex situ* U.S. and wild Namibian cheetahs from other wild populations is identified, consistent with historical records of cheetah imports. Deleterious mutations are found to be over-represented in genes associated with spermatozoal flagella in both *ex situ* and wild populations, suggesting potential causative loci of poor sperm function in cheetahs. Previously identified premature termination codons are revealed to be fixed across all cheetah populations studied. Notably, a higher proportion of unique deleterious SNPs is observed in Namibian cheetahs, highlighting important considerations for future restoration projects.

## Materials & Methods

### Samples

Whole genome re-sequencing data from 39 cheetahs was used in this study; 32 newly generated and seven from Dobrynin *et al*. (2015) (Table S*1*). Of the newly sequenced individuals, 30 were sourced from the U.S. *ex situ* population across eight institutions (Table S1). *Ex situ* samples comprised whole blood collected opportunistically in EDTA Vacutainer^®^ tubes (Becton Dickinson, U.S.) during routine veterinary exams, except for a single skeletal muscle tissue sample (AJU5640) collected during a post-mortem necropsy. The International Cheetah Studbook (L. Marker & Johnston, 2022) was used to select individuals with minimum mean kinship representing a range of modern lineages: born between 2006 and 2021, with one to nine generations of captive breeding, and a roughly equal split of males and females (16 males, 14 females; Table S1). Several known family groups were included as part of a wider study into inheritance of mutation load (Figure S*1*). The remaining two newly sequenced individuals were collected in South Sudan by Kelsey Greene at African Parks South Sudan (Table S*1*). The final dataset comprised 30 U.S. *ex situ* samples, four wild Tanzanian samples, three wild Namibian samples and two wild South Sudanese samples.

### DNA extraction and sequencing

DNA extraction, library preparation and sequencing for U.S. *ex situ* samples were conducted at Psomagen, Inc. (Maryland, U.S.) resulting in 350 bp insert libraries (see Supplementary Methods). Libraries were paired-end sequenced (2 × 150 bp) to 10x coverage on a single 10B flowcell on the Illumina NovaSeq X Plus platform. DNA extraction and library preparation for South Sudan samples were performed by Inqaba Biotec (Pretoria, South Africa) resulting in 150 bp insert libraries (see Supplementary Methods). Libraries were sequenced on a 25B flowcell on the Illumina NovaSeq X Plus platform at Admera Health (New Jersey, U.S.).

### QC, mapping and variant calling

Raw reads (fastq files) were quality checked using *FastQC* v0.11.9 (Andrews, 2010). To minimise batch effects, reads from U.S. and South Sudan samples were trimmed to 100 bp to match published samples using *Trimmomatic* v0.39 (Bolger et al., 2014). Adapters were trimmed using *Trim_galore* v0.6.10 (Krueger, n.d.). Paired and unpaired reads were mapped to the cheetah reference genome (VMU_Ajub_asm_v1.0, GCA_027475565.2, (Winter et al., 2023)) using *BWA-MEM* v0.7.17 (H. Li, 2013). Read groups were added using *GATK* v4.6.0.0 (McKenna et al., 2010) and BAM files merged and indexed with *SAMtools* v1.18 (Danecek et al., 2021).

Variant calling and initial filtering steps were performed using *GATK* v4.6.0.0 (McKenna et al., 2010) (see Supplementary Methods). Variants were filtered using default *GATK* hard filtering parameters with reduced FS (FisherStrand) requirement corresponding to our data and minimum and maximum depth of 3 (1st percentile) and 17 (99th percentile), respectively (GATK Team, 2025; Kryvokhyzha, 2016). *BCFtools* v1.22 (Danecek et al., 2021) was used to extract biallelic sites, exclude the X chromosome and exclude sites missing in > 15% of samples. Depending on the requirements of each analysis, additional filters were applied using *BCFtools* (Table *1*). For some analyses, sites with minor allele frequency (MAF) < 0.025 and related individuals (see below) were excluded. The MAF threshold was chosen to remove rare alleles likely to be sequencing errors. Linkage disequilibrium (LD) pruning was performed with r^2^ of 0.125 and window size of 50,000 kb, again only applied for some analyses (stated below). Finally, invariant sites were removed.

**Table 1.**
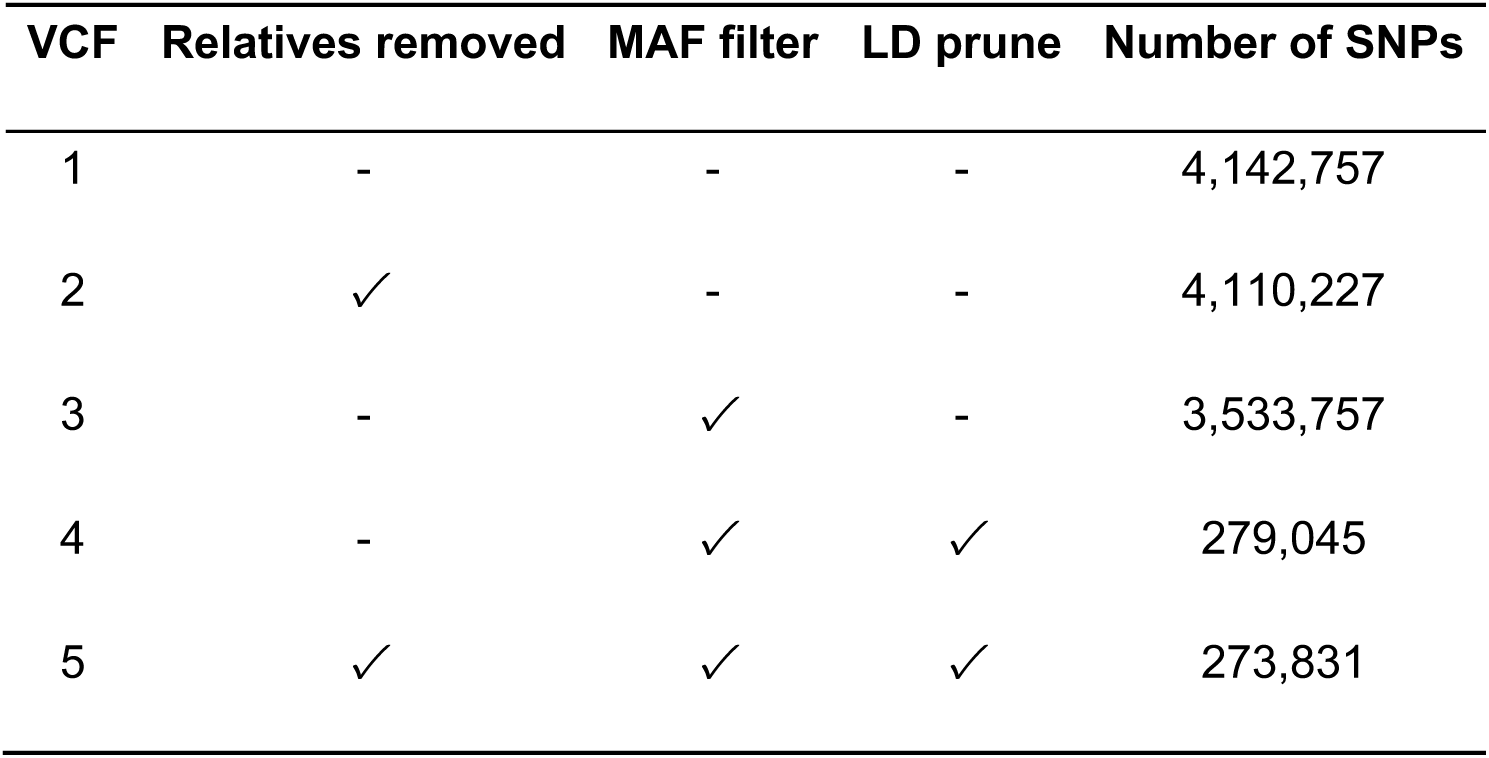
Additional filtering steps applied for each VCF file. All VCFs were filtered using GATK hard filters, biallelic sites extracted and sites missing in > 15% of samples removed. Following this, analysis-specific filters were applied. Relatives removed: related individuals (AJU10270, AJU10272, AJU9459, AJU9460, AJU9461, AJU9907, AJU9910 and AJU9890) were removed. MAF filter: sites with minor allele frequency (MAF) <0.025 were removed. LD prune: linkage disequilibrium (LD) pruning was performed with r^2^ of 0.125 and window size of 50,000 kb.

| VCF | Relatives removed | MAF filter | LD prune | Number of SNPs |
| --- | --- | --- | --- | --- |
| 1 | - | - | - | 4,142,757 |
| 2 | ✓ | - | - | 4,110,227 |
| 3 | - | ✓ | - | 3,533,757 |
| 4 | - | ✓ | ✓ | 279,045 |
| 5 | ✓ | ✓ | ✓ | 273,831 |

### Kinship

Many population genetics analyses assume unrelated individuals, kinship among our sample set was assessed to determine if any individuals needed to be removed. *PLINK* v.2.0.0 (Chang et al., 2015; Purcell et al., 2007) was used to calculate kinship using the *KING* algorithm on VCF4 (see Table *1*). Results were compared to information from the International Cheetah Studbook (L. Marker & Johnston, 2022) and confirmed expected familial relationships. Therefore, for some subsequent analyses (see Table 1), the following related individuals (offspring and full siblings) were removed from the VCF using *BCFtools* v1.22 (Danecek et al., 2021): AJU10270, AJU10272, AJU9459, AJU9460, AJU9461, AJU9907, AJU9910 and AJU9890. Where both parents were included in the analysis, all offspring were excluded. For AJU9889 and AJU9890, the individual with highest genomic coverage (AJU9889) was retained.

### Population structure and genetic diversity

To investigate population structure among U.S., Namibian, Tanzanian and South Sudanese cheetahs, a principal component analysis (PCA) was generated on VCF5 (see Table *1*) using *PLINK* v2.0.0 (Chang et al., 2015; Purcell et al., 2007) and visualised using the Python package *seaborn* (Waskom, 2021). PCAs were also run on all individuals (VCF4, see Table *1*), on the U.S. population only, and on the U.S. and Namibian individuals. *ADMIXTURE* v1.3.0 (Alexander et al., 2009) was run on VCF5 to perform ancestry estimation for K values 1-5. Pairwise *F_ST_* values were computed between populations by running *VCFtools* v0.1.16 (Danecek et al., 2011) on VCF5 in 100kb sliding windows with a 10kb step size. Mean *F_ST_* values were calculated across all windows for each pair of populations using the Weir and Cockerham estimator (Weir & Cockerham, 1984). To generate a maximum-likelihood population tree, VCF4 was converted to PHYLIP format using *vcf2phylip* v2.0 (Ortiz, 2019) and provided to *RAxML-NG* v0.8.0 (Kozlov et al., 2019). Model evaluation was performed using *RAxML-NG* v0.8.0 (Kozlov et al., 2019) to select the model with highest log likelihood, GTR+G, and the tree was generated with 1000 bootstrap replicates. A mitochondrial tree was generated as above (see Supplementary Methods).

To estimate genome-wide genetic diversity and inbreeding of each population, *VCFtools* v0.1.16 (Danecek et al., 2011) was used to calculate the inbreeding coefficient (*F_IS_*) (Weir & Cockerham, 1984; Wright, 1965), Tajima’s D (Tajima, 1989) and nucleotide diversity (π (Nei & Li, 1979)) on VCF2 (see Table *1*). Tajima’s D and π were calculated in 100 kb windows. Population-specific SNPs were identified using *BCFtools* v1.22 (Danecek et al., 2021) and plotted using R package *ggVennDiagram* (Gao et al., 2024). Observed heterozygosity (*H_O_*) was estimated for each sample using *ANGSD* v0.923 (Korneliussen et al., 2014) (see Supplementary Methods).

Runs of homozygosity (ROH) were identified from VCF3 (see Table *1*) using *RZooRoH* (Bertrand et al., 2019) in *R* v4.5.2 (R Core Team, 2025). *VCFtools* v0.1.16 (Danecek et al., 2011) was used to remove loci that were significantly out of Hardy-Weinberg Equilibrium (p-value < 0.05), as recommended by *RZooRoH* documentation. A model with multiple HBD classes was defined using the ‘zoomodel’ function, specifying seven classes with a base rate of 4, and was run on the data. The fraction of the genome in ROH (*F_ROH_*), number of ROH (*N_ROH_*), and total length of ROH (*S_ROH_*) were calculated using ROH > 1 Mb (based on the output of the Viterbi algorithm (Rabiner, 1989)) and visualized using *matplotlib* (Hunter, 2007). IDrisk (Inbreeding Depression risk), a statistic combining long ROH with non-ROH heterozygosity to predict risk of inbreeding depression, was calculated following the method by Kyriazis et al. (2025).

### Deleterious variants

*SnpEff* v5.3 (Cingolani, Platts, et al., 2012) was used to predict the functional impacts of variants in VCF1 (see Table *1*). Variants were categorised by predicted impact using *SnpSift* (Cingolani, Patel, et al., 2012). Variants predicted as high- or moderate-impact were extracted and corresponding gene names were identified. From this, *Ensembl BioMart* v115 (Cunningham et al., 2022) was used to extract corresponding human Ensembl gene IDs.

*Orthofinder* v2.5.4 (Emms et al., 2025) was run on the longest protein isoform per gene from the most recent cheetah (*A. jubatus*), human (*Homo sapiens*), dog (*Canis lupus familiaris*) and cat reference genomes (GCA_027475565.2, GCA_000001405.29, GCA_014441545.1, GCA_018350175.1, respectively). One-to-one (1:1) orthologs between the human and cheetah were identified and used as the background set for Gene Ontology (GO) enrichment analysis. The foreground set of genes were those with a corresponding 1:1 ortholog and a high or moderate impact SNP. GO enrichment analysis was run for the high and moderate impact SNPs separately, using *ShinyGO* v0.85 (Ge et al., 2020) with an enrichment false discovery rate (FDR) corrected q-value cutoff of 0.05. Population-specific SNPs with high and moderate impact were identified using *BCFtools* v1.22 (Danecek et al., 2021) and plotted using R package *ggVennDiagram* (Gao et al., 2024).

Masked and realized load for high-impact SNPs were characterised per sample by calculating the proportion of all non-missing high-impact SNPs per sample that were homozygous alternate (realized) and heterozygous (masked). The same was carried out for moderate-impact SNPs. Kruskal-Wallis and pairwise Wilcoxon rank-sum tests were carried out using SciPy Stats (Virtanen et al., 2020) and visualised with *MatPlotLib* (Hunter, 2007).

To determine the prevalence of previously identified nonsense mutations across 65 genes (Peers et al., 2025), reads for all samples were mapped to the previous cheetah reference genome, aciJub1 (GCA_001443585.1; (Dobrynin et al., 2015)) using *BWA-MEM* v0.7.17 (H. Li, 2013) (except SRR27375chewbacca, which is the sample used for this assembly). Read groups were added using *GATK* v4.6.0.0 (McKenna et al., 2010) and BAM files merged and indexed with *SAMtools* v1.18 (Danecek et al., 2021). Joint genotyping was completed using *BCFtools* v1.10.2 *mpileup* (Danecek et al., 2021) and the resultant BCF was converted to VCF format using *BCFtools view*. Filters were not applied as depth was calculated separately using *BCFtools query*. The 65 loci previously identified (Peers et al., 2025) were extracted alongside their genotype and per-sample depth using *BCFtools view* and manually examined to determine prevalence in the population data.

## Results

### QC, mapping and variant calling

The total number of paired-end reads per individual ranged from 103,469,302 to 494,748,773 with a mean depth of 4.2 to 18.5x (Table S2). Mapping efficiency to the reference genome was high, with > 98.9% of reads successfully aligned per individual. 6,896,511 variants were called across 39 individuals. From this, 4,986,770 SNPs were extracted. After *GATK* quality filtering, 4,269,772 SNPs remained, 4,210,600 of which were biallelic. Further filtering of biallelic SNPs removed 873,535 sites due to missing genotype data and 1,032,070 sites due to the minor allele threshold, leaving 2,304,995 SNPs. LD pruning removed 1,862,015 sites, leaving 442,980 SNPs. Eight related individuals were removed, leaving 31 individuals.

### Kinship

Pairwise kinship coefficients were calculated between all individuals and visualised as heatmaps (Figures 1, S2). Individuals within the U.S. *ex situ* population had higher kinship compared to the other populations, although Namibian individuals also showed weakly positive kinship to the U.S. population (Figure *S2*). In comparison, South Sudanese and Tanzanian cheetahs showed lower kinship. Within the *ex situ* U.S. population (Figure *1*), pairwise kinship values were generally low, but elevated kinship was observed within known family groups identified in the International Cheetah Studbook. For example, kinship values of 0.25 (suggesting first-degree relatives) were calculated between AJU10270 and AJU10272, as well as between these individuals and AJU7225 and AJU8965. This is consistent with the studbook record listing AJU10270 and AJU10272 as offspring of AJU7225 and AJU8965.

**Figure 1.**
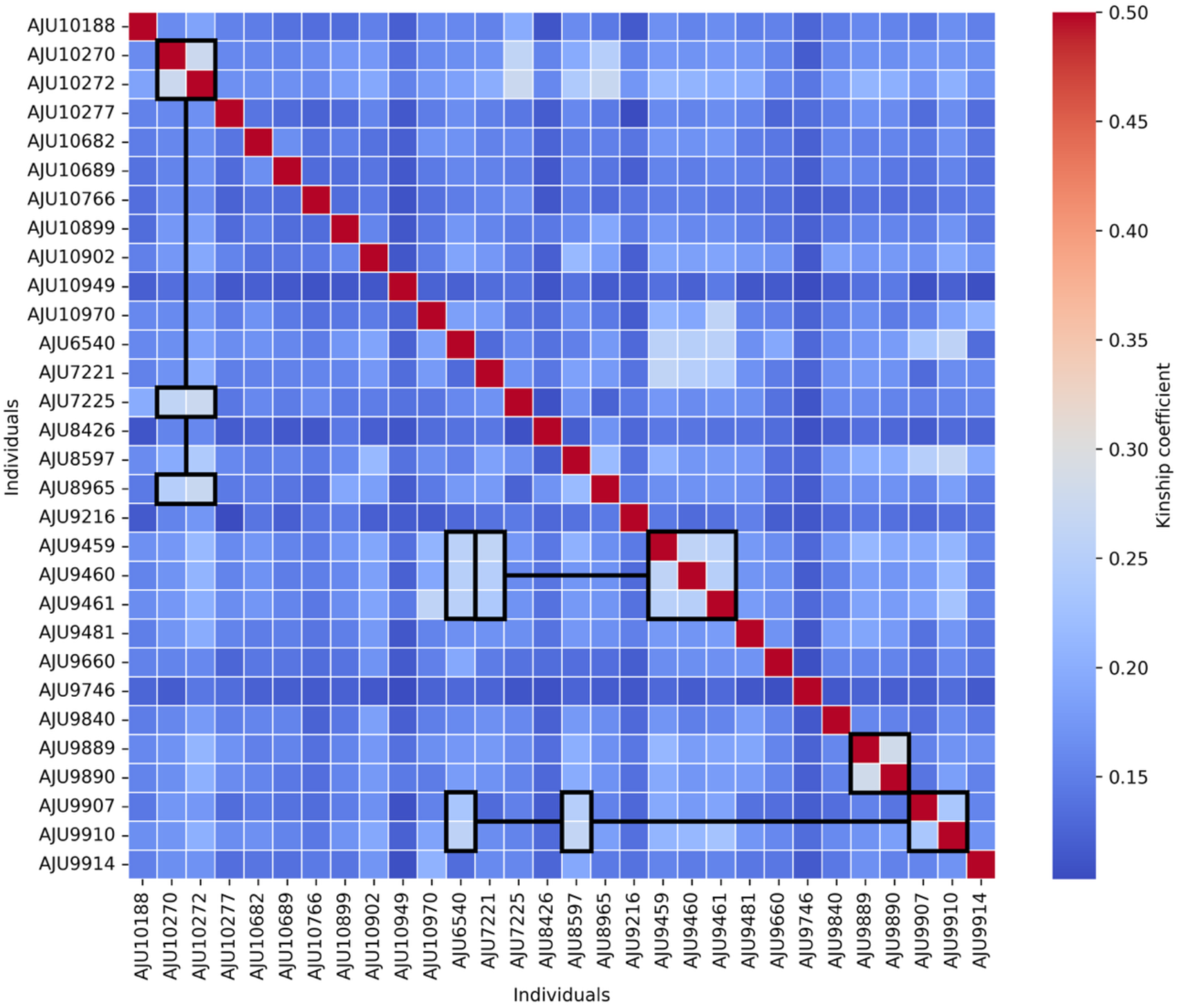
Pairwise kinship values for *ex situ* U.S. cheetahs. Each cell represents the estimated kinship coefficient between two individuals, with red indicating higher relatedness and blue indicating negative values, suggesting unrelatedness. The diagonal represents self-relatedness (kinship coefficient = 0.5). Known family groups based on the studbook (see Figure S1) are shown with black boxes and lines connecting offspring and parents. Kinship between most pairs of individuals is low, but first degree family relationships can be observed between known family groups.

### Population structure and genetic diversity

Distinct separation was identified between South Sudan, Tanzanian, and Namibian and *ex situ* U.S. cheetahs (Figure *2A*). PC1, separating cheetahs from South Sudan and Tanzania from those in Namibia and the U.S., represented 12.91% of diversity. PC2, which shows variation between the African populations, represented 8.25% of diversity, with remaining PCs representing less than 6% each (Figure S3). A PCA with all individuals, including relatives, showed a consistent pattern (Figure S4). PCAs of the U.S. and Namibian populations (Figure S5,6) showed separation of Namibian individuals from the U.S. population, however the variation between U.S. and Namibian individuals was less than that observed within the U.S. population.

**Figure 2.**
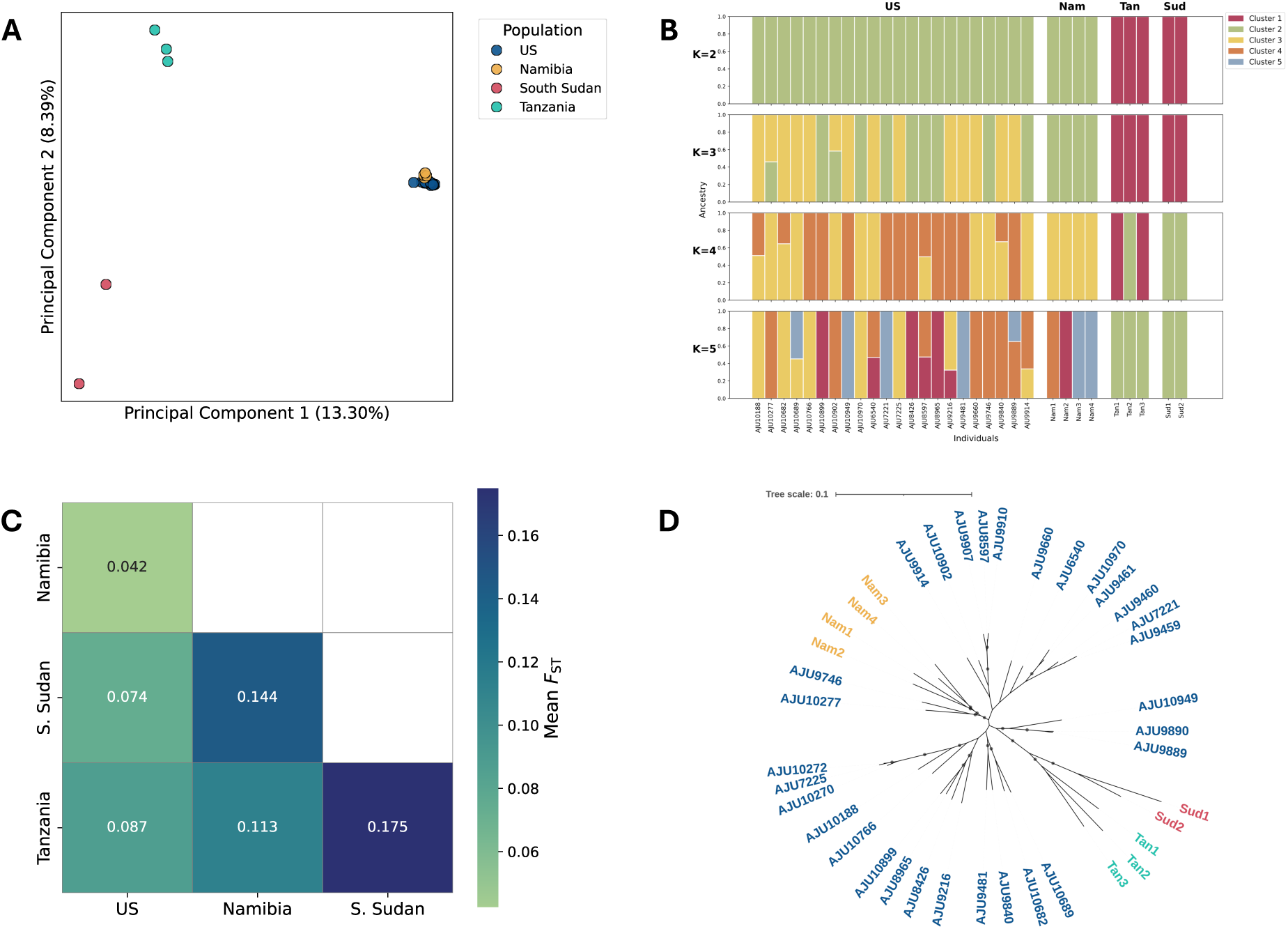
Population structure of *ex situ* (U.S.) and wild (Namibia, South Sudan, Tanzania) cheetahs. (**A**) **Principal Component Analysis (PCA) of unrelated cheetahs.** PCA based on filtered genome-wide SNPs showing genetic clustering by population: U.S. (blue), Namibia (yellow), South Sudan (red), and Tanzania (green). The percentage of variance explained by each principal component is shown in axes labels. (**B**) **ADMIXTURE analysis of unrelated cheetahs.** Model-based clustering of cheetah populations at K = 2–5 using only unrelated individuals. Each vertical bar represents an individual, and colours represent the proportion of ancestry assigned to each genetic cluster. Admixture cross-validation errors for K = 1 to 5 were 0.54, 0.60, 0.69, 0.89 and 0.89, respectively. (**C**) **Pairwise genetic differentiation between cheetah populations.** Heatmap showing mean pairwise Weir and Cockerham’s *F_ST_* values among cheetah populations. Darker colours represent greater genetic differentiation, with lighter colours indicating higher genetic similarity. (**D**) **Maximum likelihood phylogenetic tree of cheetah samples based on genome-wide SNPs.** Sample colours match plot A. The scale bar indicates substitutions per site. Branches with bootstrap values > 90% are marked with a grey circle.

Admixture cross validation (CV) errors for K = 1 to 5 were 0.54, 0.60, 0.69, 0.89 and 0.89, respectively with the lowest error at K = 1, suggesting no population structure (Figure *2B*; *S7*). Genetic differentiation between populations, indicated by pairwise *F_ST_* values (Figure *2C*), mirrored the pattern observed in the PCA (Figure *2A*). The U.S. and Namibian populations showed the lowest differentiation (*F_ST_* = 0.042), suggesting substantial shared ancestry. Greatest differentiation was observed between South Sudan and Tanzania (*F_ST_* = 0.175), suggesting strong genetic divergence. The population phylogeny based on genome-wide SNPs mirrored the population structure observed in the PCA and ADMIXTURE analyses, with clear separation of the South Sudan and Tanzanian cheetahs from the U.S. and Namibian individuals (Figure *2D*). Bootstrap support was high (> 90) for most branches, particularly those separating South Sudan from Tanzania and the Namibian individuals from those in the U.S. The mitogenome tree showed South Sudanese and Tanzanian cheetahs clustered within the U.S. population, although branch lengths were very low (Figure S8).

Population genetic diversity measures showed higher inbreeding and lower genetic diversity in the South Sudan population (Figure *3*). *F_IS_* was comparatively lower in the U.S. and Namibian populations (Figure 3A); however, this result was not significant (Kruskal-Wallis: H=0.65, p=0.88). Observed heterozygosity (*H_O_*) differed significantly across populations (Kruskal-Wallis: H=20.29, p=1.48x10^-4^; Figure 3B), with a global mean of 0.000682 and a range of 0.000498 to 0.000940. *H_O_* was highest in South Sudan and lowest in Namibia and Tanzania, although these pairwise differences were not statistically significant (Bonferroni-corrected Wilcoxon rank sum, p > 0.05), likely due to small sample size (n ≤ 4). Tajima’s D was positive in all populations, consistent with demographic contraction, and was highest in the U.S. population, potentially indicative of a founder effect or population contraction associated with captivity (Figure 3C). A significant overall difference was identified (Kruskal-Wallis: H=5919.46, p=0) and pairwise Wilcoxon rank-sum tests identified a significant difference between all pairs of populations (Table S3). Nucleotide diversity (π) was also significantly different between populations (Kruskal-Wallis: H=4486.73, p=0), with significant differences between all pairs of populations except the U.S. and Tanzania (Figure 3D; Table S3). Median π ranged from 0.00032 to 0.00037.

**Figure 3.**
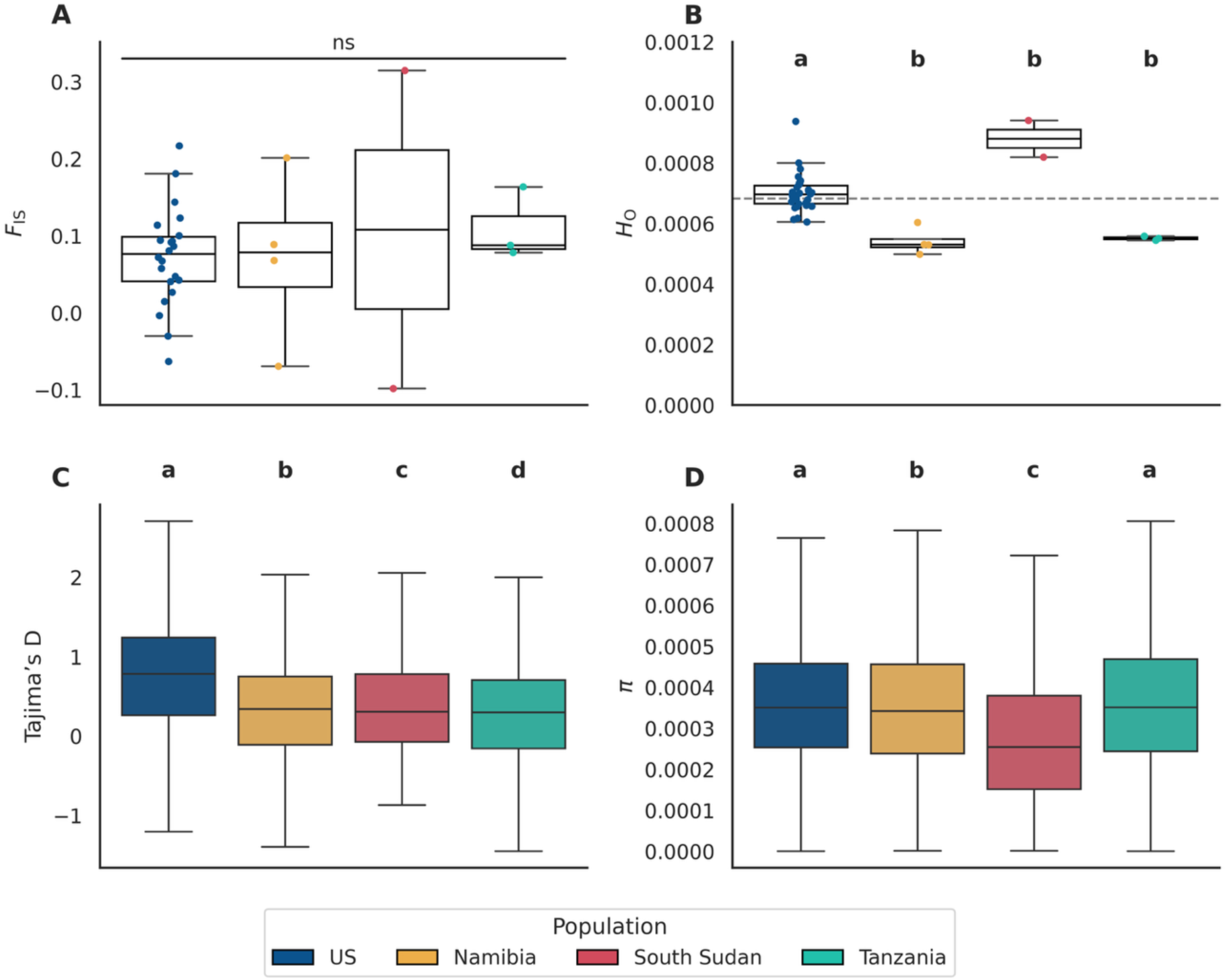
Genetic diversity statistics across cheetah populations. Boxplots showing population-level estimates of (**A**) inbreeding coefficient (*F_IS_*), (**B**) observed heterozygosity (*H_O_*), (**C**) Tajima’s D (100 kb windows), and (**D**) nucleotide diversity (π, 100 kb windows). For all plots, colours indicate populations: U.S. (blue), Namibia (yellow), South Sudan (red), and Tanzania (green), and significant results (Pairwise Wilcoxon rank-sum tests and Bonferroni correction, p > 0.05) are indicated by labels (a, b, c, d) for each plot. See Table S3 for full significance values. In plot **B**, a dashed line shows the mean *H_O_* across all populations.

ROH suggested the highest inbreeding in the South Sudan population (Figure *4*), with highest *F_ROH>1Mb_* and longer ROH compared to the other populations. The South Sudan and Tanzanian populations display a larger proportion of ROH in higher HBD classes, suggesting greater historic inbreeding in these populations (Figure 4A). There were generally fewer and shorter ROH in the US population, with several outliers. Individuals AJU9216 and AJU7225 had higher *N_ROH>1Mb_* and *S_ROH>1Mb_* than the rest of the population and a higher proportion of ROH in lower homozygous-by-descent (HBD) classes, suggesting more recent inbreeding. Sample AJU7225 showed the highest *F_ROH>1Mb_* in the dataset, with a notably higher proportion of ROH in low HBD classes. When visualising the distribution of ROH across the genome (Figure 4D, E), long stretches of ROH can be observed in sample AJU7225 but not in its offspring (AJU10270). Although a Kruskal–Wallis test indicated an overall difference in *F_ROH>1Mb_* among populations (H = 10.73, p = 0.013), no individual pairwise comparison was significant after Bonferroni correction (Wilcoxon rank-sum tests: all adjusted p > 0.05). *F_ROH>1Mb_* ranged from 0.014 to 0.211, with a global *F_ROH>1Mb_* mean of 0.048. IDrisk scores showed low risk of population inbreeding depression in all populations, based on the score thresholds provided by (Kyriazis et al., 2025) (Figure S9).

**Figure 4.**
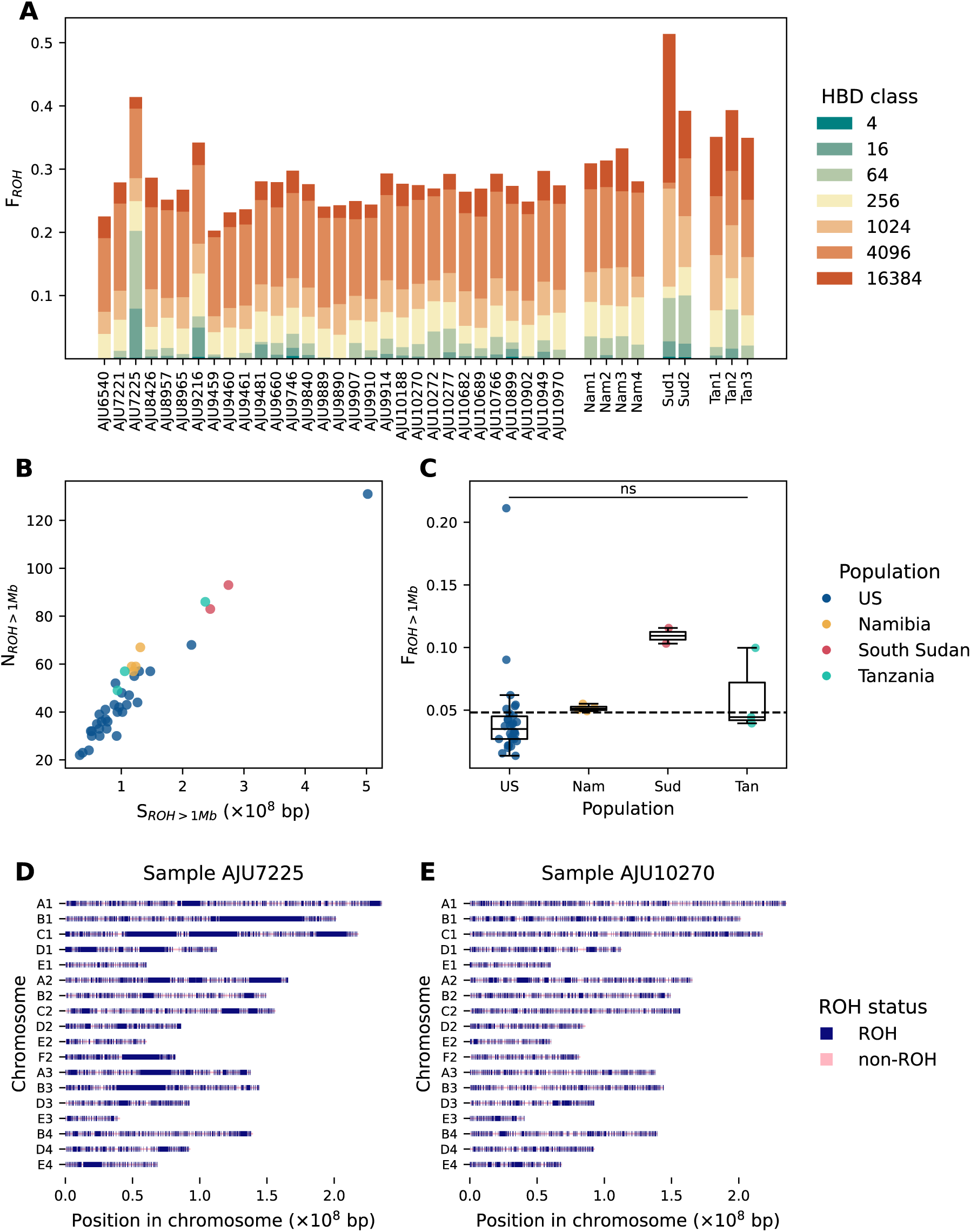
Runs of homozygosity (ROH) across cheetah populations. (**A**) **ROH partitioned into homozygosity-by-descent (HBD) classes for each individual.** HBD classes reflect approximately twice the number of generations since inbreeding, with more recent ROH in green and more ancient ROH in red. (**B**) **The relationship between the number of ROH segments (*N_ROH>1Mb_*) and the sum of ROH lengths (*S_ROH>1Mb_*),** where colours correspond to populations: U.S. (blue), Namibia (yellow), South Sudan (red), and Tanzania (green). (**C**) **Fraction of the genome in ROH >1Mb (*F_ROH>1Mb_*) per population**, with a data point for each individual overlaid on the boxplot. Colours correspond to populations as shown in (C). Pairwise Wilcoxon rank-sum tests found no significance (all Bonferroni-corrected p-values >0.05), indicated by ’ns’ label. (**D**) **Distribution of ROH across the genome of AJU7225**, where non-ROH is shown in light pink and ROH is shown in dark blue. (**E**) **Distribution of ROH across the genome of AJU10270**, offspring of AJU7225. Colours correspond to those in plot D.

### Deleterious variants

Population-specific biallelic SNPs were identified in all populations, with most SNPs unique to the U.S. population as expected based on sample size (Figure *S10*). The Namibian population had the fewest unique SNPs, with almost 95% of Namibian SNPs also identified in the U.S. population. Deleterious variants were identified using SnpEff, classifying 684 variants as high impact, 18,448 as moderate impact, 27,506 as low impact and 4,142,676 as modifiers (variant is outside coding regions or its impact is unknown).

703 annotated genes were associated with the 684 high impact SNPs. Of these, 562 had a 1:1 ortholog in the human and so were used as the foreground in GO enrichment analysis. Eleven Biological Process GO terms were significantly (FDR-corrected q < 0.05) overrepresented in this gene list, mostly related to cilium and sperm flagellum (Figure *5*; Table *S5*).

**Figure 5.**
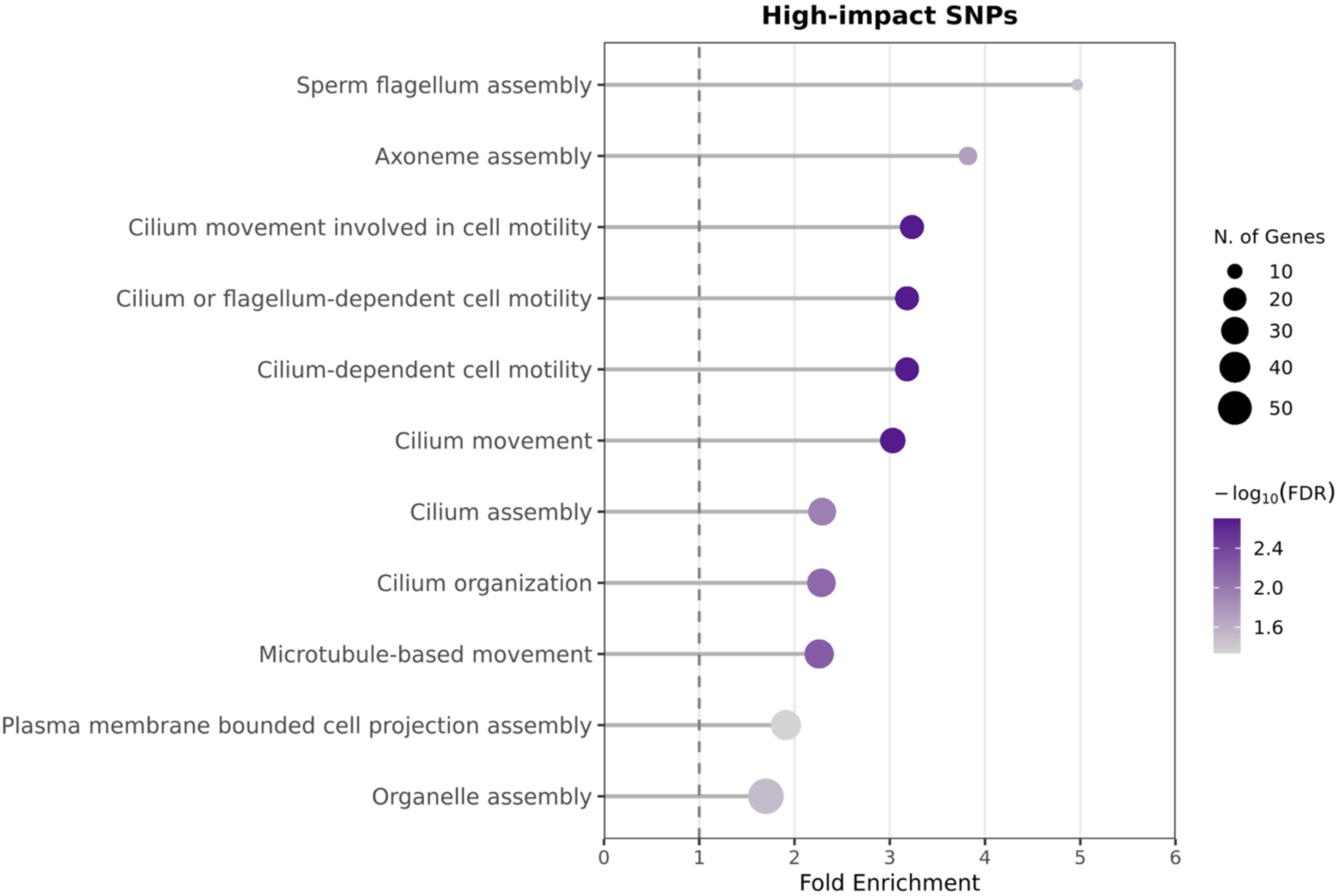
Gene ontology (GO) enrichment of genes with predicted high impact SNPs in cheetahs. GO Biological Process terms significantly enriched (FDR-corrected q-value < 0.05) among genes containing high-impact SNPs across the four cheetah populations. Fold enrichment is shown on the x-axis, with the size of the point reflecting the number of genes annotated with each GO term. The colour of each point shows the statistical significance (-log_10_ FDR) of the enrichment of each GO term, where darker points are more significantly enriched. A fold enrichment score of 1, shown by a dotted line, means no enrichment in the gene set.

The same analysis was then completed for SNPs with moderate impact. 7,818 genes were associated with the 18,448 moderate impact SNPs. Of these, 6365 had a 1:1 ortholog in the human; this set of genes was used as the foreground for this GO enrichment analysis. 143 Biological Process GO terms were significantly enriched (FDR-corrected q < 0.05) in this gene set. The top 20 most significant (Figure *6*; Table S6) were mostly involved in cilium movement and microtubules.

**Figure 6.**
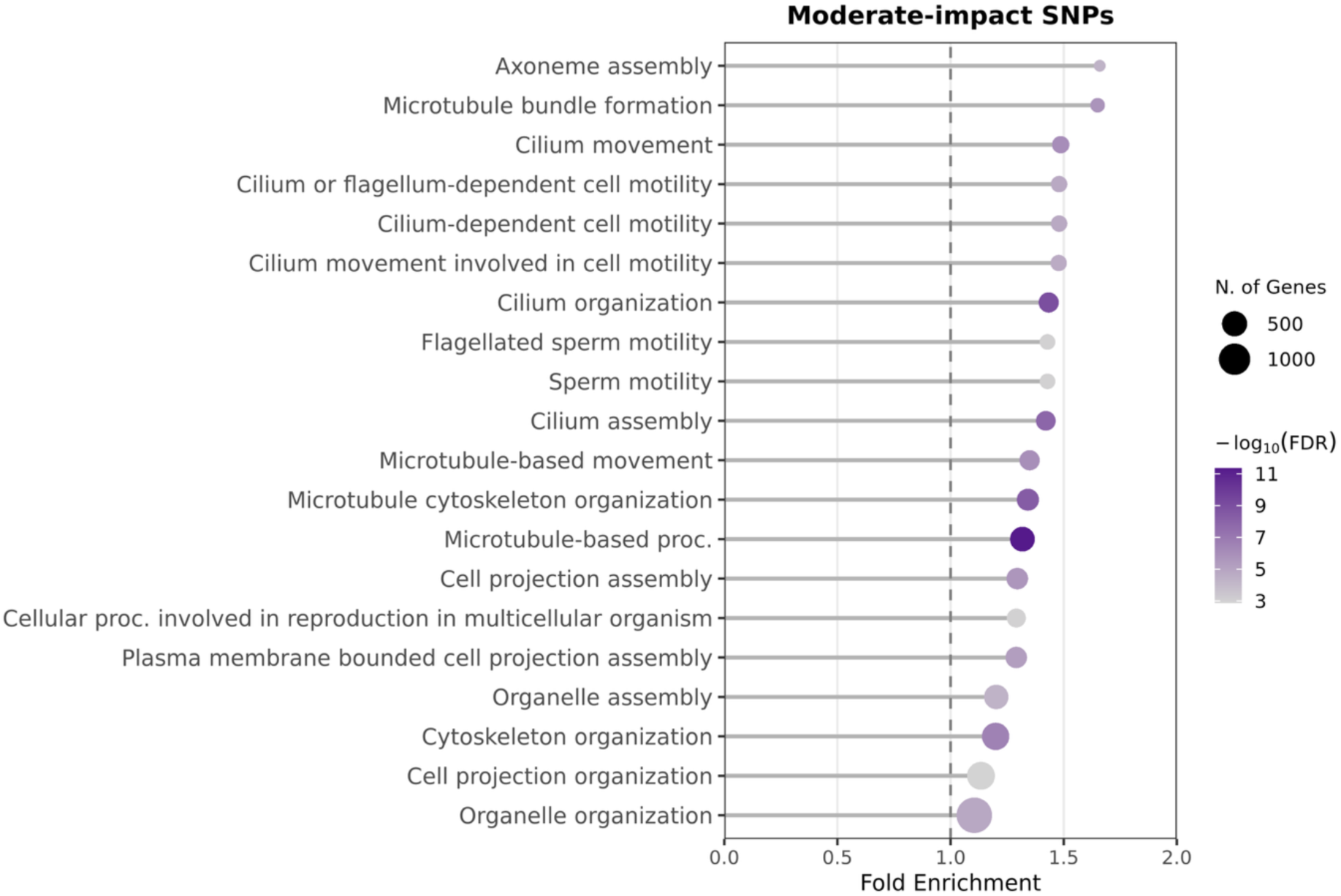
Gene ontology (GO) enrichment of genes with predicted moderate impact SNPs in cheetahs. GO Biological Process terms significantly enriched (FDR-corrected q-value < 0.05) among genes containing moderate-impact SNPs across the four cheetah populations. Fold enrichment is shown on the x-axis, point size corresponds to the number of genes annotated with each GO term and the colour of each point shows the statistical significance (-log_10_ FDR) of the enrichment, where darker points are more significantly enriched. A fold enrichment score of 1, shown by a dotted line, means no enrichment in the gene set.

High and moderate impact SNPs mirrored that of the total SNP set across the four populations, with each population containing at least 41 unique SNPs with predicted high impact and at least 528 unique SNPs with predicted moderate impact (Figure *7*). When comparing these values to overall numbers of population-specific SNPs, the Namibian population contained a higher proportion of unique high-impact SNPs than other populations, whilst Tanzania contained a lower proportion of unique high- and medium-impact SNPs.

**Figure 7.**
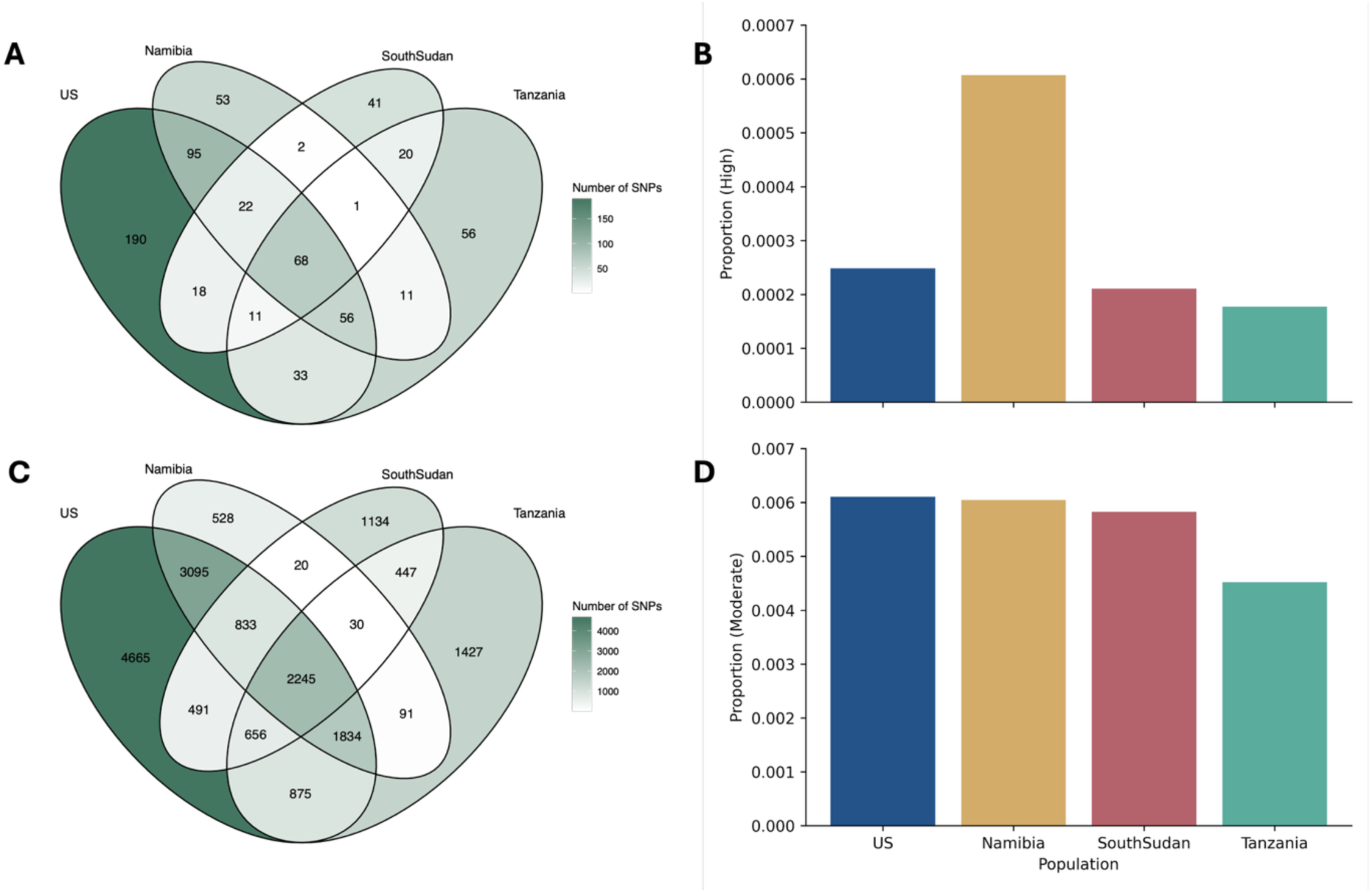
Distribution of predicted high- and moderate-impact SNPs among cheetah populations. (A) Distribution of high-impact SNPs predicted by SnpEff between the four populations. The number of SNPs specific to each combination of populations is shown in each segment, with a darker colour indicating a higher number of SNPs. (**B**) Unique high-impact SNPs as a proportion of the overall number of unique SNPs per population (see Figure 8). Panels **C** and **D** show the same information as panels A and B, respectively, for moderate-impact SNPs.

The majority of high-impact SNPs had a low allele frequency (Figure S11), with similar patterns observed in moderate- and low-impact SNPs (Figures S12, S13). Sixty-four high-impact SNPs with an allele frequency over 0.9 were extracted and their distribution across populations was plotted (Figure S11B). A large proportion of high-frequency SNPs unique to South Sudan was observed, likely due to the small sample size. Forty unique genes were associated with these SNPs. GO enrichment analysis found no significant enrichment (FDR-corrected q > 0.05) of these genes with a high-frequency, high-impact SNP.

Masked and realized load were assessed per sample as a proportion of total SNPs (Figure 8). A significant overall difference was identified for realized load of both high- and moderate-impact SNPs (Kruskal-Wallis, high: H=12.9344, p=0.00478; moderate: H=15.7594, p=0.00127) and for moderate-impact masked load (Kruskal-Wallis: H=7.85295, p=0.0491), but not for high-impact masked load (Kruskal-Wallis: H=0.165128, p=0.983). Bonferroni-corrected pairwise Wilcoxon rank-sum tests identified significant differences between both high- and moderate-impact realized load between the US and South Sudan and the US and Tanzania, as well as moderate-impact masked load between the US and Tanzania (Table S4; Bonferroni-corrected p < 0.05). Realized load was highest in South Sudan and Tanzania for both masked and realized load, with a roughly equal proportion of masked load across populations.

**Figure 8.**
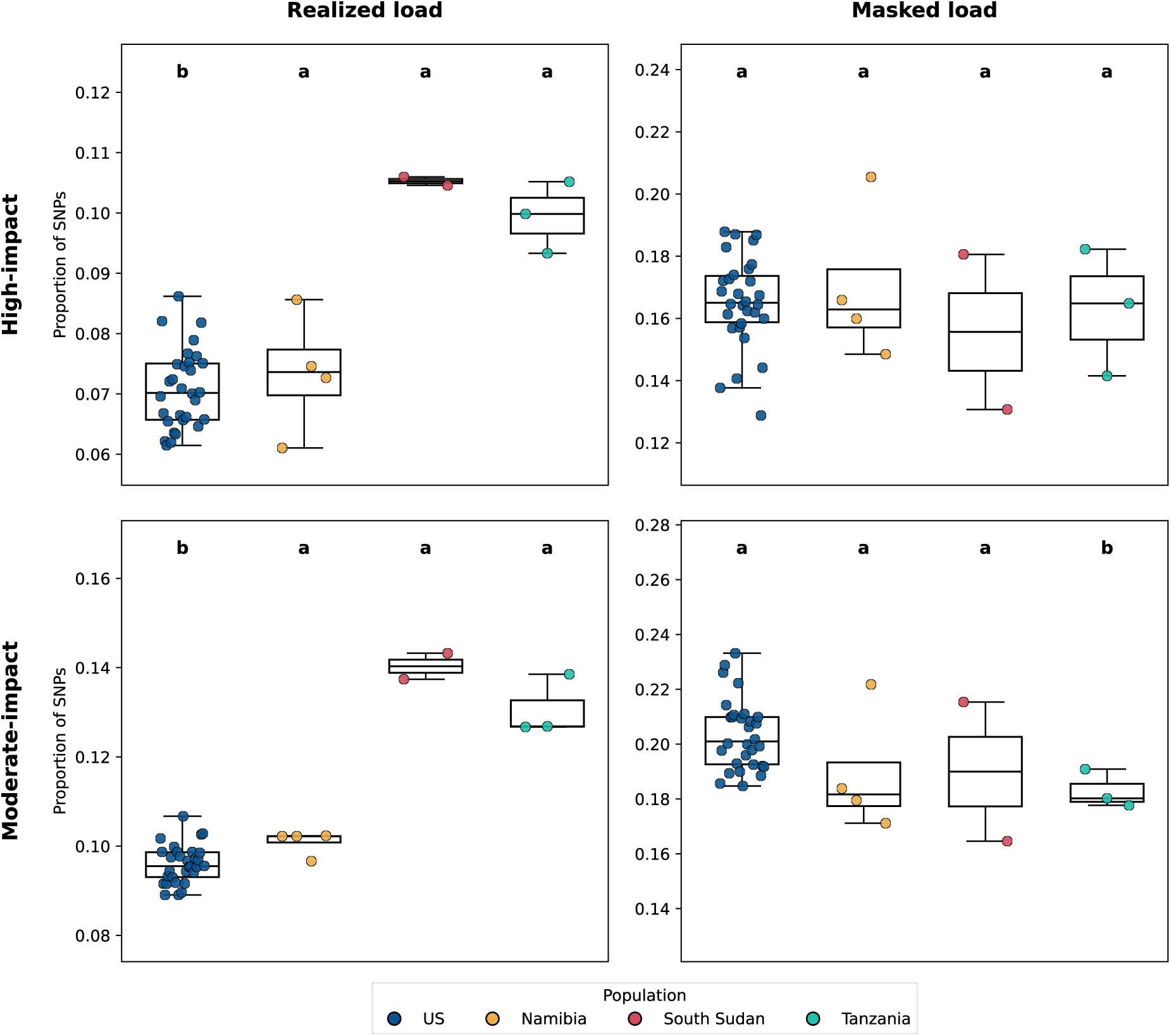
Realized and masked load of high- and moderate-impact SNPs as predicted by SnpEff. For all plots, colours indicate populations: U.S. (blue), Namibia (yellow), South Sudan (red), and Tanzania (green), and significant results (Pairwise Wilcoxon rank-sum tests and Bonferroni correction, p > 0.05) are indicated by labels (a, b, c, d) for each plot. See Table S4 for full significance values. (A) Proportion of homozygous high-impact SNPs out of the total number of high-impact SNPs per sample. (B) Proportion of heterozygous high-impact SNPs out of the total number of high-impact SNPs per sample. (C) Proportion of homozygous moderate-impact SNPs out of the total number of moderate-impact SNPs per sample. (D) Proportion of heterozygous moderate-impact SNPs out of the total number of moderate-impact SNPs per sample.

Of the 65 previously identified premature termination codons (PTCs) (Peers et al., 2025), 36 were not found in any population data and were therefore considered unique to the reference genome (aciJub1, GCA_001443585.1, (Dobrynin et al., 2015)) as was previously reported ((Peers et al., 2025), Table S7). Two loci did not have sufficient coverage to determine population distribution. Twenty-three loci were homozygous for the reference allele in all individuals, meaning the PTC-causing mutation is fixed across these samples. The remaining four loci had some variation in the population data. Two were indels, suggesting a false SNP call in the previous analysis. One mutation, in the Four And A Half LIM Domains 5 gene (*FHL5*), was only found in eight *ex situ* U.S. samples and was otherwise not present in the population data. The final mutation, in the Defensin Beta 116 gene (*DEFB116*), was fixed in all but two *ex situ* samples, a mother and daughter (AJU8965 and AJU10270).

## Discussion

Understanding the genetic diversity and distribution of masked and realised deleterious mutations across cheetah populations provides insight into the severity of past bottlenecks and their contemporary impact on population health. Here, we present the first whole genome analysis of genetic health of an *ex situ* cheetah population relative to wild cheetahs. We identify a significant over-representation of deleterious mutations in sperm-associated genes, particularly in Namibian cheetahs, whilst Tanzanian cheetahs contained the lowest proportion of deleterious SNPs. We show low genetic differentiation between *ex situ* and wild Namibian cheetahs, consistent with the proposed origin of the majority of *ex situ* cheetahs, and comparative levels of genetic diversity between *ex situ* and wild populations.

### Population structure and relatedness

The genomic evidence presented here corroborates data in the International Cheetah Studbook, as genomic kinship values reflect the familial relationships recorded in the studbook. This is an important result, as *ex situ* breeding pairs are usually selected based on estimates of minimal kinship from the studbook, which relies on decades of diligent record keeping. Therefore, ensuring that the studbook is accurate is crucial to prevent accidental inbreeding in *ex situ* conditions (Ballou et al., 2010; Galla et al., 2022).

Population structure analyses consistently showed *ex situ* U.S. and wild Namibian cheetahs clustering together, reflecting the origin of *ex situ* cheetahs in the U.S (Marker-Kraus, 1997). A few individuals from eastern African populations were brought into captivity prior to the 1960s (Marker-Kraus, 1997), which may explain the lower genetic differentiation between South Sudan and Tanzanian populations with the U.S. compared to wild-wild differentiation. We observed lower *F_ST_* values across all populations compared to a previous study based on double-digest restriction site associated DNA (ddRAD) sequencing, despite similar numbers of samples used (Prost et al., 2022). This discrepancy likely stems from the varying resolution of the data types; whole-genome data can produce more robust estimates of *F_ST_* than reduced-representation approaches such as ddRAD sequencing (Lou et al., 2021).

Namibian and Tanzanian cheetahs are currently recognised as the same subspecies (*A. j. jubatus*) by the IUCN, although the Tanzanian population was previously classified as *A. j. raineyi* (Krausman & Morales, 2005). Based on *F_ST_* estimations, the Namibian and Tanzanian samples used in this study show proportionally high genetic differentiation, supporting (Prost et al., 2022)’s call for the IUCN to return to the previous subspecies classification. However, as only four Namibian and three Tanzanian cheetahs were included here, a larger sample set is needed to confirm this observation. Additionally, admixture analysis found no significant separation between samples.

### Genetic diversity and measures of inbreeding

Across all sampled cheetah populations, measures of genetic diversity were low, as has previously been observed in the cheetah (Castro-Prieto et al., 2011; Dobrynin et al., 2015; O’Brien et al., 1983; Terrell et al., 2016). Nucleotide diversity (π) values were comparable to previous studies (Dobrynin et al., 2015); median π values were between 0.00032-0.00039, supporting the idea first proposed by (O’Brien et al., 1983) that cheetahs contain very low genetic diversity. π was significantly lower in South Sudan and Namibia than *ex situ* US cheetahs, supporting a previous observation of continued decline in genetic diversity in wild cheetahs, although this pattern was not observed in Tanzania (Terrell et al., 2016). Contrasting π, observed heterozygosity (*H_O_*) was highest in South Sudan, although this result was not significant and is likely skewed by small sample size (Nazareno et al., 2017). All four populations contained higher *H_O_* than previous studies, with a global mean of 0.000682, compared to previous estimations of 0.000400 and 0.000440 (Kim et al., 2016; Prost et al., 2022). Despite this difference, our results support previous observations that cheetahs contain low heterozygosity compared to other mammals (Morin et al., 2021; Pečnerová et al., 2021; Prost et al., 2022). Additionally, positive Tajima’s D values support the theory that all populations have experienced a genetic bottleneck or population contraction (Dobrynin et al., 2015; Fabiano et al., 2025; Menotti-Raymond & O’Brien, 1993).

Measures of inbreeding were high across all populations, again supporting previous research (Dobrynin et al., 2015; Menotti-Raymond & O’Brien, 1993; O’Brien & Johnson, 2005; Terrell et al., 2016). Median *F_IS_* was greater than zero in all populations, suggesting all populations have experienced inbreeding. The lack of significant difference in inbreeding between wild and *ex situ* cheetahs also supports previous observations (Prost et al., 2022). However, as with the previous study, sample sizes for wild cheetah populations remain low, meaning more extensive sampling is required to confirm this finding. *F_ROH_* (the fraction of the genome in ROH above a specified length) has often been used to compare inbreeding between populations (McQuillan et al., 2008). Here, we demonstrate that *F_ROH>1Mb_* does not significantly differ between cheetah populations. When comparing the distribution of ROH across the genome to previous studies in the cheetah, we observe far less homozygosity, likely due to differences in the assembly quality of the reference genome (contig N50 = 35.1 Kbp versus 96.8 Mbp for aciJub1 and VMU_Ajub_asm_v1.0 assemblies, respectively) and methodology used to call ROH (Ceballos et al., 2018; Shafer & Kardos, 2025; Shi et al., 2026). This is because a more contiguous genome assembly significantly increases accuracy when detecting different size classes of ROH, particularly for longer ROH segments (Shi et al., 2026). Additionally, we observe lower *F_ROH>1Mb_* than many other wild and *ex situ* mammal populations, including those with historic inbreeding and low effective population sizes, suggesting that cheetahs may exhibit a less homozygous genome than was previously suggested (Armstrong et al., 2024; Brüniche-Olsen et al., 2018; Dussex et al., 2021; Khan et al., 2021; Leon-Apodaca et al., 2023; Niehaus et al., 2025; Quinn et al., 2024; Sánchez-Barreiro et al., 2021).

Although no significant difference in *F_ROH>1Mb_* was observed between populations, South Sudanese cheetahs had higher *N_ROH>1Mb_* and *S_ROH>1Mb_* than the majority of US and all Namibian and Tanzanian individuals, suggesting more recent inbreeding in this population. A greater proportion of ROH in high HBD classes was also observed in the South Sudan population, suggesting this population experienced greater historic inbreeding than the other populations. This population also had a higher proportion of ROH in low HBD classes, suggesting more recent inbreeding, although the highest proportion of ROH in the lowest HBD classes was observed in US sample AJU7225. This individual contains a greater number of ROH and total length of ROH, resulting in the highest *F_ROH>1Mb_* in the dataset. This suggests significant recent inbreeding in this individual, which was confirmed by its pedigree, which shows several recent cases of inbreeding between half-siblings (Figure *S14*). However, as two of AJU7225’s offspring were included in this study, it is possible to show that by mating AJU7225 with an individual with lower *F_ROH>1Mb_*, their offspring can be prevented from inheriting the long ROH observed in their father (e.g., Galla et al., 2020). This finding highlights that recent inbreeding can be reverted provided effective action is taken.

Our data show that the *ex situ* U.S. cheetah population exhibits comparable measures of inbreeding and genetic diversity to the wild populations, suggesting *ex situ* breeding programmes are effectively managed to prevent genetic diversity loss, as has been previously suggested (Williams & Hoffman, 2009; Witzenberger & Hochkirch, 2011). This is strong support for the cheetah *ex situ* breeding programme, as reduced genetic diversity and increased mutational load (see next section) due to a captivity-induced founder effect are not observed. With effective management, *ex situ* populations can act as important reservoirs of genetic diversity for the future.

### Deleterious variants

SNPs with a predicted ’high’ or ’moderate’ deleterious impact within protein-coding genes were significantly enriched for functions associated with male reproductive fitness. Although the relationship between inbreeding and male fertility in cheetahs has been widely debated (Crosier et al., 2018; Fitzpatrick & Evans, 2009; Terrell et al., 2016; Wildt et al., 1983), this finding provides strong evidence that cheetahs have experienced an accumulation of deleterious mutations in sperm-associated genes. This corroborates previous research focusing on premature termination codons and human-cheetah orthologs of fertility-related genes, where highly deleterious mutations were identified in several sperm-associated genes (Dobrynin et al., 2015; Peers et al., 2025). Despite observing high levels of ROH and a relatively high *F_IS_* in Tanzania compared to the other populations, this population contains the lowest proportion of unique high- and moderate-impact SNPs. Despite this, realised load (homozygous high- and moderate- impact SNPs) is highest in Tanzania and South Sudan, although this result is not significant. This pattern of high realized load is expected in South Sudan, which had higher *F_ROH_* and therefore higher homozygosity across the genome. This observation may be skewed by low sample size, so future work should aim to sample more individuals from this population. It is also worth noting that SNPs are not polarised, so ancestral and derived alleles are not separated.

When comparing the distribution of these high- and moderate-impact SNPs across populations, we observe a significantly higher proportion of unique high-impact SNPs related to male fertility in Namibian individuals, although realized load (homozygous deleterious SNPs) is low in Namibia compared to the other wild populations. Comparison of semen samples across wild populations are crucial to assess the impacts of such potentially deleterious SNPs, as previous studies of wild cheetah semen have focused on the southern African subspecies (Crosier et al., 2007; Lindburg et al., 1993; Wildt et al., 1983, 1988). As the majority of *ex situ* U.S. cheetahs are sourced from the Namibian population, the difference in the proportion of high-impact, deleterious SNPs between Namibian and *ex situ* individuals could indicate that these mutations arose more recently in the Namibian population, or that they have been removed from the *ex situ* population through purging or controlled breeding. However, as only four genomes from Namibian cheetahs were used in this study, future work to sequence more genomes from this population is necessary to confirm this finding.

It is important to note that the SNP counts of population-specific mutations are affected by the reference genome used to call SNPs, raising the important issue of reference bias in population genetic studies, which can underestimate genetic diversity and differentiation (Akopyan et al., 2025). As the reference genome used here was generated from an *ex situ* cheetah housed at Lisbon Zoo, Portugal, we would expect this bias to falsely reduce the number of unique SNPs in the *ex situ* U.S. cheetahs and inflate the number of unique SNPs found in South Sudan and Tanzania. Despite this, the U.S. cheetahs still have a higher proportion of unique high and moderate impact SNPs than South Sudan and Tanzania, suggesting that reference bias has not impacted the overall patterns we observe.

Our findings have significant implications for cheetah conservation management programmes and integrated conservation strategies (One Plan Approach). Understanding the genomic divergence between wild and *ex situ* populations is critical for assessing the suitability of captive-bred individuals for future reintroduction or translocation efforts. Augmentation, a form of translocation whereby individuals are moved between subpopulations to maintain or bolster genetic diversity and reduce inbreeding, has previously been applied to cheetahs in southern Africa (Buk et al., 2018; L. L. Marker et al., 2008). Confirming whether the elevated levels of highly deleterious SNPs we observe in Namibia are consistently distributed across the southern African population is necessary to assess any potential threat of facilitating the spread of such mutations. Cheetahs from southern Africa are also being relocated as part of efforts to reintroduce the species to India (Tordiffe et al., 2023; Venugopal, 2025). Sourcing individuals from a population with a high mutation load and exposing them to a founder effect could result in these deleterious SNPs becoming exposed through increased homozygosity (Kyriazis et al., 2021; Robinson et al., 2023). Again, further sampling across the southern African population is necessary to confirm our observation of high mutation load and inform ongoing and future translocation programmes.

Previous research has shown that genomics-informed captive breeding can reduce inbreeding and mutational load, such as in the pink pigeon (*Nesoenas mayeri*) where genetic load was used to identify optimal mate pairs with minimal load in offspring (Speak et al., 2024). In the pink pigeon, this method is more feasible due to the low number of individuals in captivity, making it possible to sample and sequence whole genomes of a larger proportion of the *ex situ* population. In species with larger *ex situ* populations, like the cheetah, the time and cost of collecting and sequencing every breeding individual in captivity makes this method infeasible. However, identification of deleterious mutations, particularly those impacting key functions like male fertility, segregating at high frequency in the *ex situ* population could enable the generation of a target-capture SNP panel that could be used to sequence these loci and prevent fixation of such deleterious mutations through mutational load-informed breeding recommendations (Bertola et al., 2022; Kleinman-Ruiz et al., 2017; Wehrenberg et al., 2024).

In conclusion, through resequencing of *ex situ* and wild cheetah populations, we present the first whole genome comparison of *ex situ* and wild cheetah populations. We observe low genetic diversity across populations but a more heterozygous genome than other inbred species. We confirm the fixation of PTCs in multiple genes associated with male fertility across *ex situ* and wild cheetahs and identify an over-representation of deleterious SNPs across sperm-associated genes, with a high load of these unique to Namibia. Finally, we show that the *ex situ* cheetah population holds comparable genetic diversity and lower measures of inbreeding compared to wild populations and highlight an empirical example of the use of outbreeding to prevent long ROH being passed to offspring. Our results emphasise the importance of integrating genomic data to inform conservation management of threatened species. Such data are critical for maintaining the adaptive potential of *ex situ* populations and ensuring that captive-bred individuals remain genetically compatible with their wild counterparts for future reintroduction.

## Acknowledgements

We thank Kelsey Greene, Jordana Morgan, and colleagues at African Parks South Sudan for sample collection and provision from South Sudan. We are also grateful to the following institutions for providing samples from the United States: White Oak Conservation, Smithsonian’s National Zoo and Conservation Biology Institute, San Diego Zoo Wildlife Alliance, Fossil Rim Wildlife Center, Wildlife Safari, Fort Worth Zoo, Toronto Zoo, and Toledo Zoo. We also thank Dr Graham Etherington (Earlham Institute, UK) for providing the gatk_params.py script.

## Data Accessibility and Benefit-Sharing statement

Raw sequence reads are available in the European Nucleotide Archive: PRJEB110820. Benefits from this research arise from the sharing of our data on public databases as described above. A collaboration was developed with conservation organisations from South Sudan from which we obtained genetic samples and our research addresses some of their concerns regarding the conservation management of the cheetah.

## Author Contributions

Language used to describe roles below uses the CRediT Taxonomy (credit.niso.org).

Conceptualization: WH, KPK

data curation: JAP, HS

formal analysis: JAP, HS

funding acquisition: KPH, EEA, WH

investigation: JAP, HS

methodology: JAP, HS, WH, KPH, WJN

project administration: JP, HS, EEA, AEC, WJN, KPK, WH

resources: EEA, AEC, WJN, KPK, WH

software: JP, HS, EEA, WH

supervision: WH, KPK, WJN

validation: JP, HS

visualization: JP, HS

writing—original draft: JP

writing—review & editing: JP, HS, EEA, AEC, WJN, KPK, WH

## Ethics approval statement

Samples for this study were collected opportunistically (e.g. during routine veterinary practices or necropsies). All *ex situ* samples were provided with full permission from each zoological facility. This project complied with all Convention on Biological Diversity and the Convention on International Trade in Endangered Species of Wild Fauna and Flora (CBD and CITES) regulations.

