## Supplementary Information for "Over-representation of sperm-associated deleterious mutations across wild and *ex situ* cheetah (*Acinonyx jubatus*) populations"

**Affiliations:**

**Table of contents**

| Supplementary Methods | 1 |
| --- | --- |
| Supplementary Figures | 3 |
| Supplementary Table legends | 13 |

Supplementary Methods

**DNA extraction**

DNA extraction, library preparation and sequencing for U.S. *ex situ* samples were conducted at Psomagen, Inc. (Maryland, U.S.). Genomic DNA extraction was conducted using the Mag-Bind Blood and Tissue Kit (Omega Bio-Tek Inc., Georgia, U.S.). DNA concentration and quality were assessed using PicoGreen fluorometry (Victor X2; Life Technologies, California, U.S.), an Agilent 4200 TapeStation (Agilent Technologies, California, U.S.), and 1% agarose gel electrophoresis. Genomic libraries were prepared using the TruSeq DNA PCR Free Library Preparation Kit (Illumina, California, U.S.), with an insert size of 350 bp following enrichment of this fragment length via ultrasonication using a Covaris S220 Ultrasonicator (Covaris, Massachusetts, U.S.). Sample AJU6540 required a second DNA extraction and library preparation due to low initial DNA quantity. Libraries were quality-checked and quantified using an Agilent 4200 TapeStation and LightCycler qPCR assay (Roche Life Science, Missouri, U.S.) then paired-end sequenced (2 × 150 bp) to 10x coverage on a single 10B flowcell on the Illumina NovaSeq X Plus platform. Raw reads were then demultiplexed by Psomagen, Inc.

For wild South Sudan samples, DNA extraction and library preparation was performed by Inqaba Biotec (Pretoria, South Africa). Extractions were performed using the Quick-DNA Miniprep Plus Kit (Zymo Research, U.S.) and library preparation was conducted using the NEBNext Ultra II Library Preparation Kit (New England Biolabs, Massachusetts, U.S.), generating 150 bp insert libraries. Libraries were sequenced on a 25B flowcell on the Illumina NovaSeqX Plus platform at Admera Health (New Jersey, U.S.).

**Variant filtering**

Variant calling and initial filtering steps were performed using *GATK* v4.6.0.0 (McKenna et al., 2010). For each sample, the workflow included the following steps: *SortSam*, *MarkDuplicates*, and *HaplotypeCaller* (run in ploidy-aware mode for male samples), followed by execution of the gatk_params.py script ([*https://github.com/TGAC/JP_PhD/blob/main/gatk_params.py*](https://github.com/TGAC/JP_PhD/blob/main/gatk_params.py)). Variants were processed through *VariantFiltration* and *SelectVariants*, after which *BaseRecalibration* and *ApplyBQSR* were applied. *HaplotypeCaller* was then run again to produce the recalibrated variant calls. GVCF files were indexed using *IndexFeatureFile* and a database of variants was generated using *GenomicsDBImport*. Finally, a multisample VCF was created using this database with *GenotypeGVCFs*. Single Nucleotide Polymorphisms (SNPs) were extracted using *SelectVariants* and variants were filtered using default *GATK* hard filtering parameters with reduced FS (FisherStrand) requirement corresponding to our data and minimum and maximum depth of 3 (1st percentile) and 17 (99th percentile), respectively (GATK Team, 2025; Kryvokhyzha, 2016).

**Mitochondrial tree**

To generate a mitochondrial tree, mitogenomes were assembled from the FASTQ files for all individuals. *Cutadapt* v4.7 (Martin, 2011) was used to trim Illumina sequencing adapters from all reads. To identify reads corresponding to nuclear pseudogenes of mitochondrial origin (NUMTs), *Numt Parser* (de Flamingh et al., 2023) was run on each trimmed FASTQ file. Since no NUMT reference was available for the cheetah, we used identified NUMTs and a mitochondrial reference genome (NC_016470.1) from the puma, *Puma concolor* (Biró et al., 2025), which is positioned within the same felid subclade as the cheetah (G. Li et al., 2016). Cheetah FASTQ files were aligned to NUMTs and the mitochondrial genome with *Bowtie2* v2.5.3 (Langmead & Salzberg, 2012) in fast-local mode. *Numt Parser* identifies reads as “numt”, “cymt” (cytoplasmic mitochondrial origin), or “undetermined”; reads identified as “cymt” were filtered from the FASTQ files using *seqkit* v2.8.1 (Shen et al., 2016) and *seqtk* v1.4 (H. Li, 2023).

A cheetah mitochondrial genome reference sequence (CM050222.1 (Winter et al., 2023)) was annotated with *MITOS2* v2.1.10 (Donath et al., 2019). The sequence between genes trnP and trnF (bases 15445-16986) was removed, deleting the control region. The script “repair.sh” from *bbmap* v39.06 (Bushnell, 2014) was used to fix read pairing in the mitochondrial FASTQ files, which were aligned to the edited reference sequence with *Bowtie2* v2.5.3 (Langmead & Salzberg, 2012) in fast-local mode. SAM files were sorted and consensus sequences (including deletions) were extracted with *SAMtools* v1.19.2. Consensus sequences were aligned with *MAFFT* v7.525 (Katoh & Standley, 2013) using default settings. A tree was generated from this alignment using *RAxML-NG* v0.8.0 (Kozlov et al., 2019). Model evaluation was performed using *RAxML-NG* v0.8.0 (Kozlov et al., 2019) to select the model with highest log likelihood, GTR+G, and the tree was generated with 1000 bootstrap replicates.

**Observed heterozygosity**

Observed heterozygosity (*H_O_*) was estimated for each sample BAM file using *ANGSD* v0.923 (Korneliussen et al., 2014) (see Supplementary Materials). For each sample, we generated a per-individual site allele frequency (SAF) spectrum with genotype likelihood model 2 and a minimum base quality of 30, as per (Pečnerová et al., 2021) to enable comparison to previously published estimations. Because ANGSD requires an ancestral sequence for SAF generation, we supplied the reference genome as the ancestral sequence and estimated a folded spectrum. We then inferred the one-dimensional site frequency spectrum (1d-SFS) for each individual using *realSFS* (default settings). Individual *H_O_* was calculated as the proportion of heterozygous loci in the per-individual 1d-SFS.

Supplementary Figures

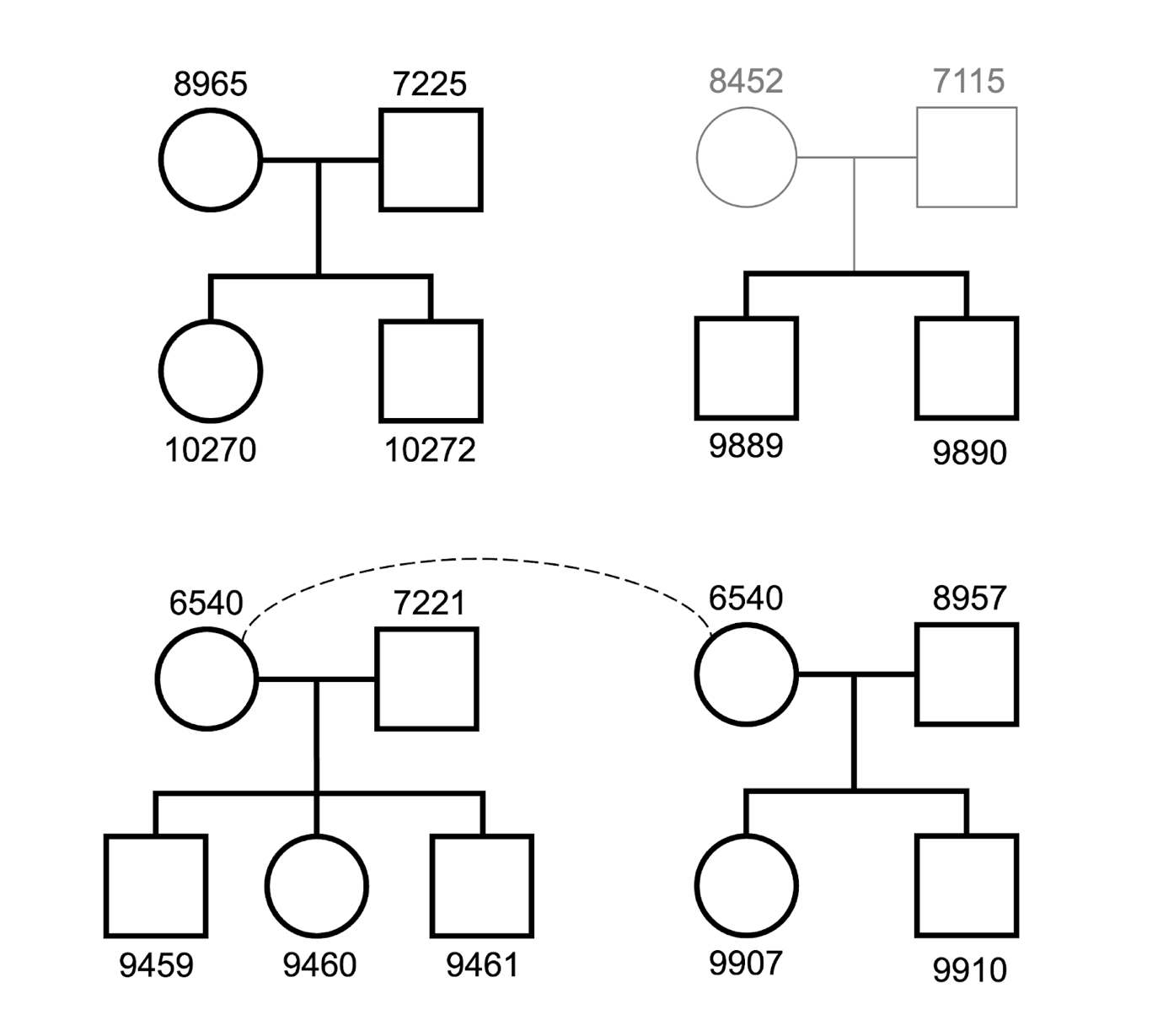

**Figure S1. Pedigrees of cheetah family groups included in this study.** Females are represented as circles and males as squares. Individuals shaded in grey (AJU8452, AJU7115) were not sampled, but their offspring were included. The dashed line indicates that female AJU6540 contributed offspring to two different family groups.

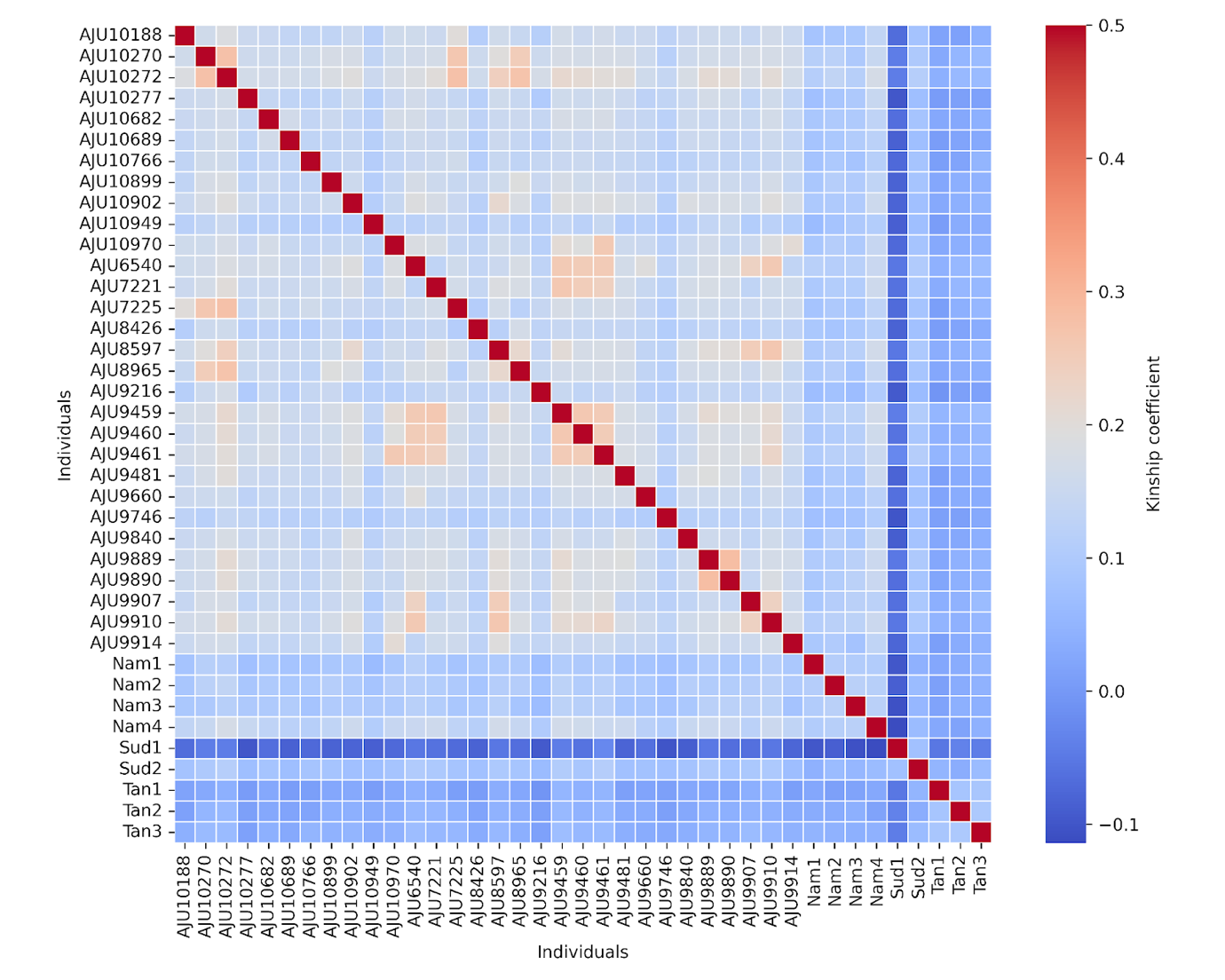

**Figure S2. Pairwise kinship values for all cheetahs in this study.** Each cell represents the estimated kinship coefficient between two individuals, with red indicating higher relatedness and blue indicating negative values, which suggest unrelatedness. The diagonal represents self-relatedness (kinship coefficient = 0.5). High kinship is observed within the U.S., Namibia and Tanzania populations and U.S. and Namibian cheetahs share weakly positive kinship. Negative kinship is observed between South Sudan and Tanzanian populations compared to U.S. and Namibian cheetahs.

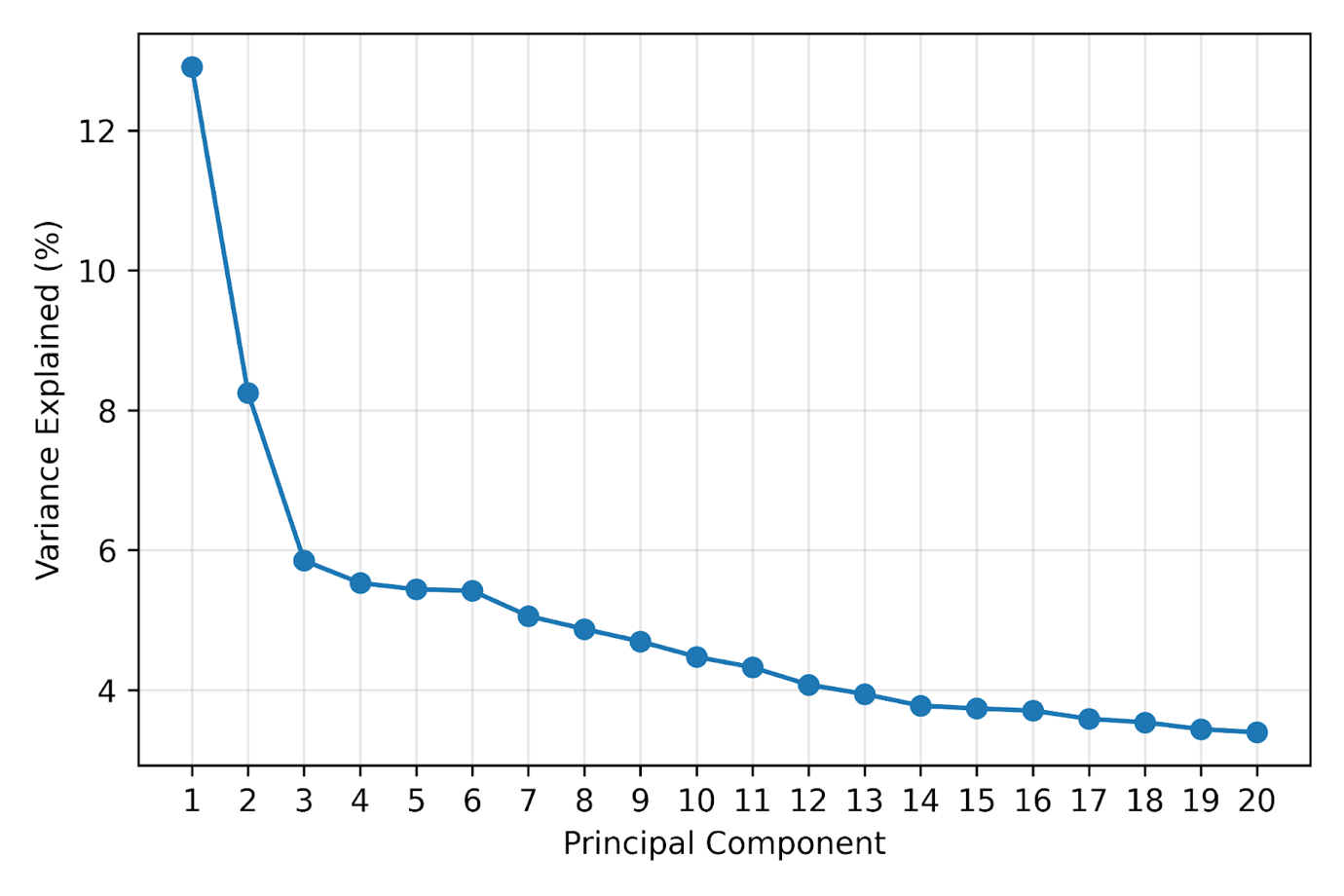

**Figure S3. Scree plot showing variance explained by each principal component of a principal component analysis (PCA) run on unrelated cheetahs (Figure 4).** Principal components 1 and 2 explain at least 8% of variance each, with subsequent PCs each contributing to < 6% each.

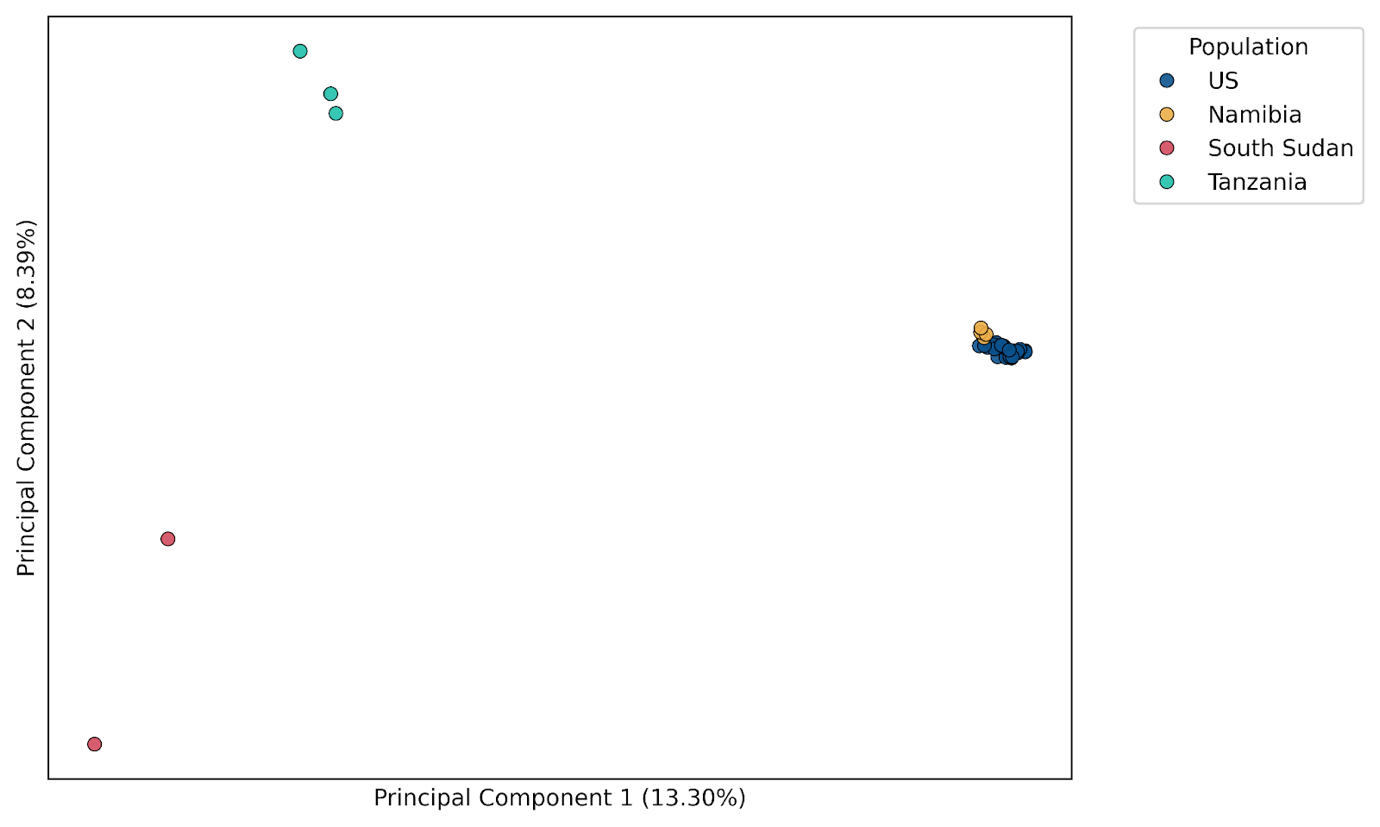

**Figure S4. Principal Component Analysis (PCA) of all 39 cheetahs used in this study.** PCA based on filtered genome-wide SNPs showing genetic clustering by population: U.S. (blue), Namibia (yellow), South Sudan (red), and Tanzania (green). The percentage of variance explained by each principal component is shown in axes labels.

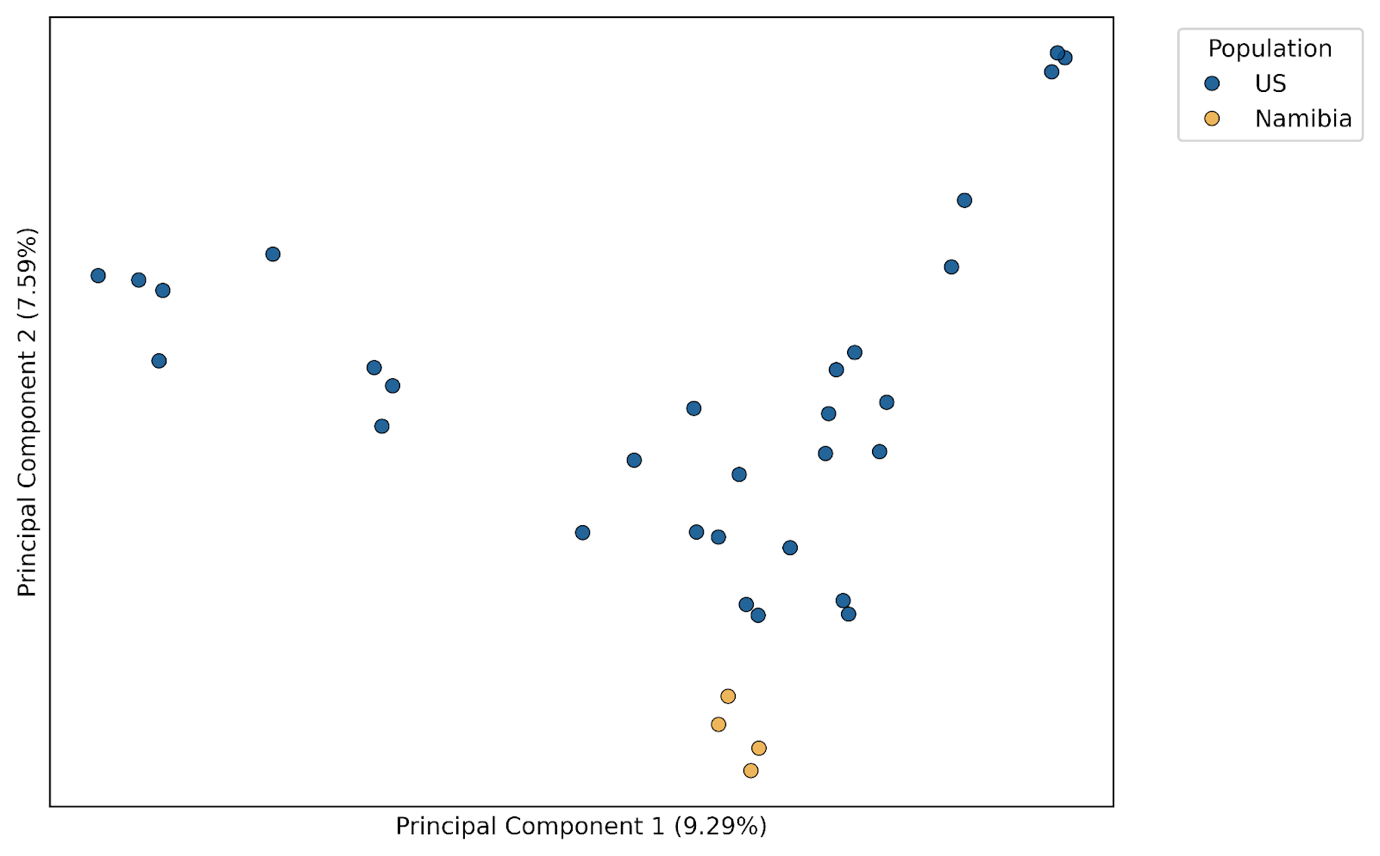

**Figure S5. Principal Component Analysis (PCA) of U.S. and Namibian cheetahs.** PCA based on filtered genome-wide SNPs of captive U.S. (blue) and wild Namibian (yellow) cheetahs. The percentage of variance explained by each principal component is shown in axes labels.

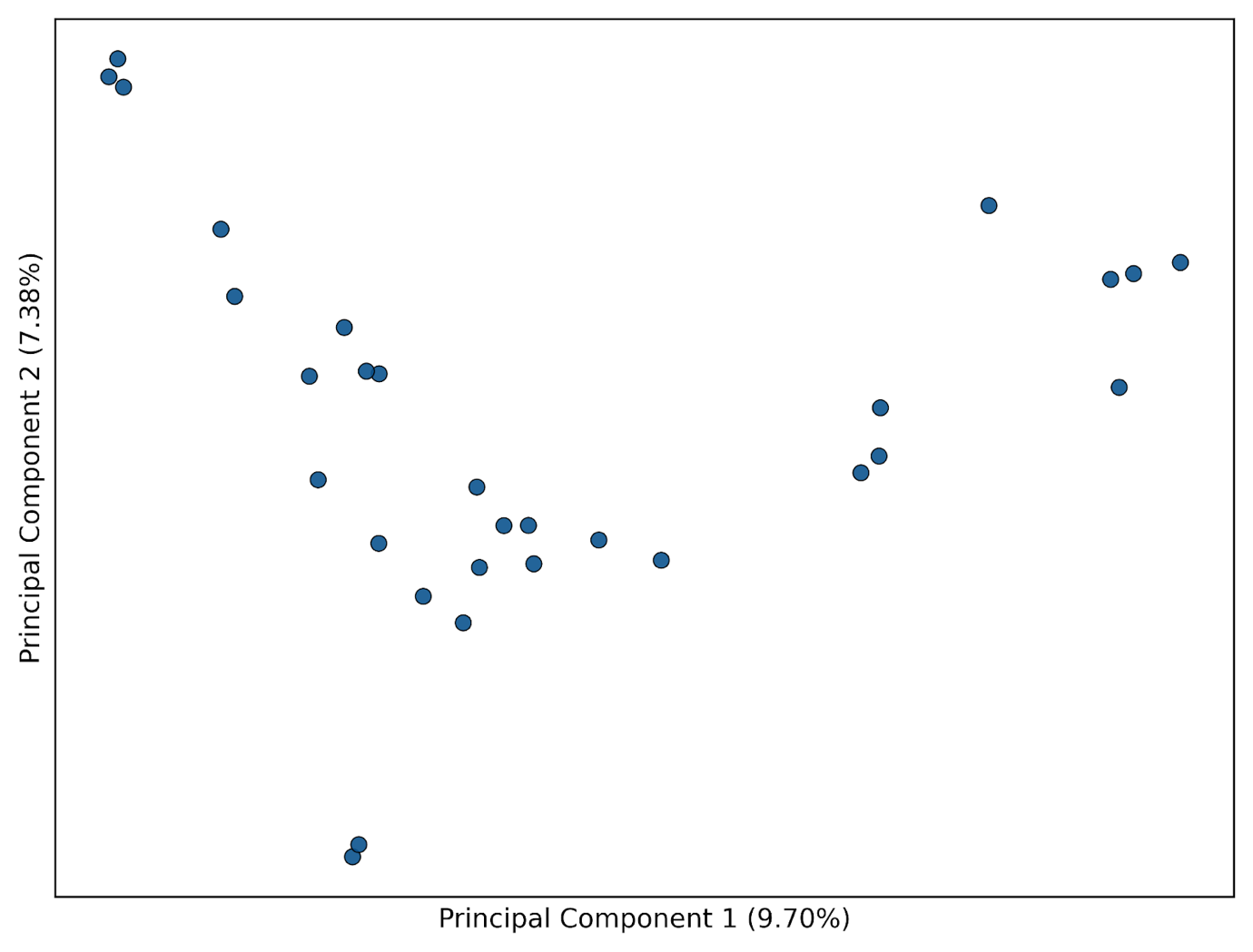

**Figure S6. Principal Component Analysis (PCA) of captive U.S. cheetahs.** PCA based on filtered genome-wide SNPs of US cheetahs. The percentage of variance explained by each principal component is shown in axes labels.

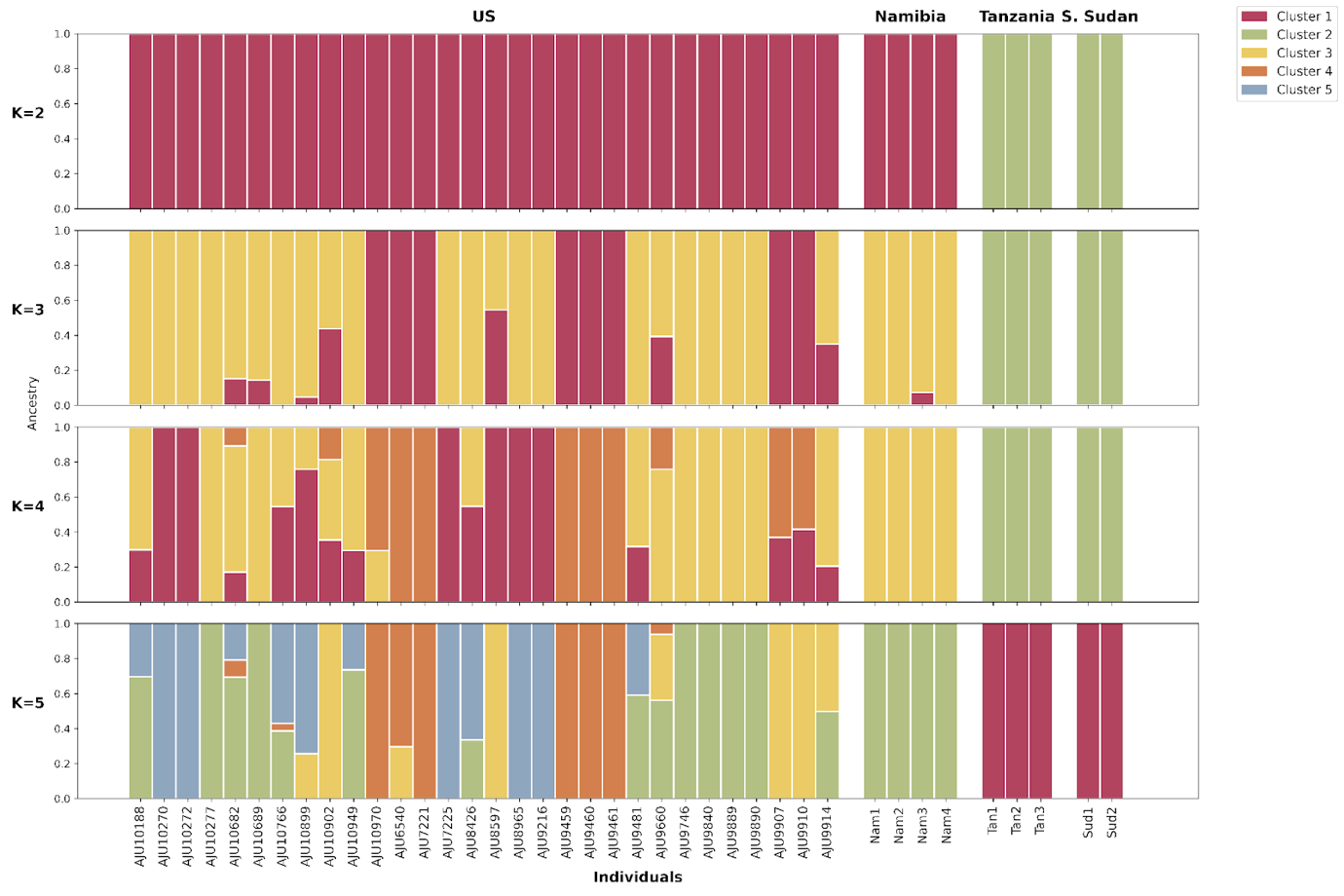

**Figure S7. ADMIXTURE analysis of all cheetahs in this study.** Model-based clustering of cheetah populations at K = 2–5 using only unrelated individuals. Each vertical bar represents an individual, and colours represent the proportion of ancestry assigned to each genetic cluster. Admixture cross-validation errors for K = 1 to 5 were 0.49, 0.52, 0.55, 0.57 and 0.62, respectively.

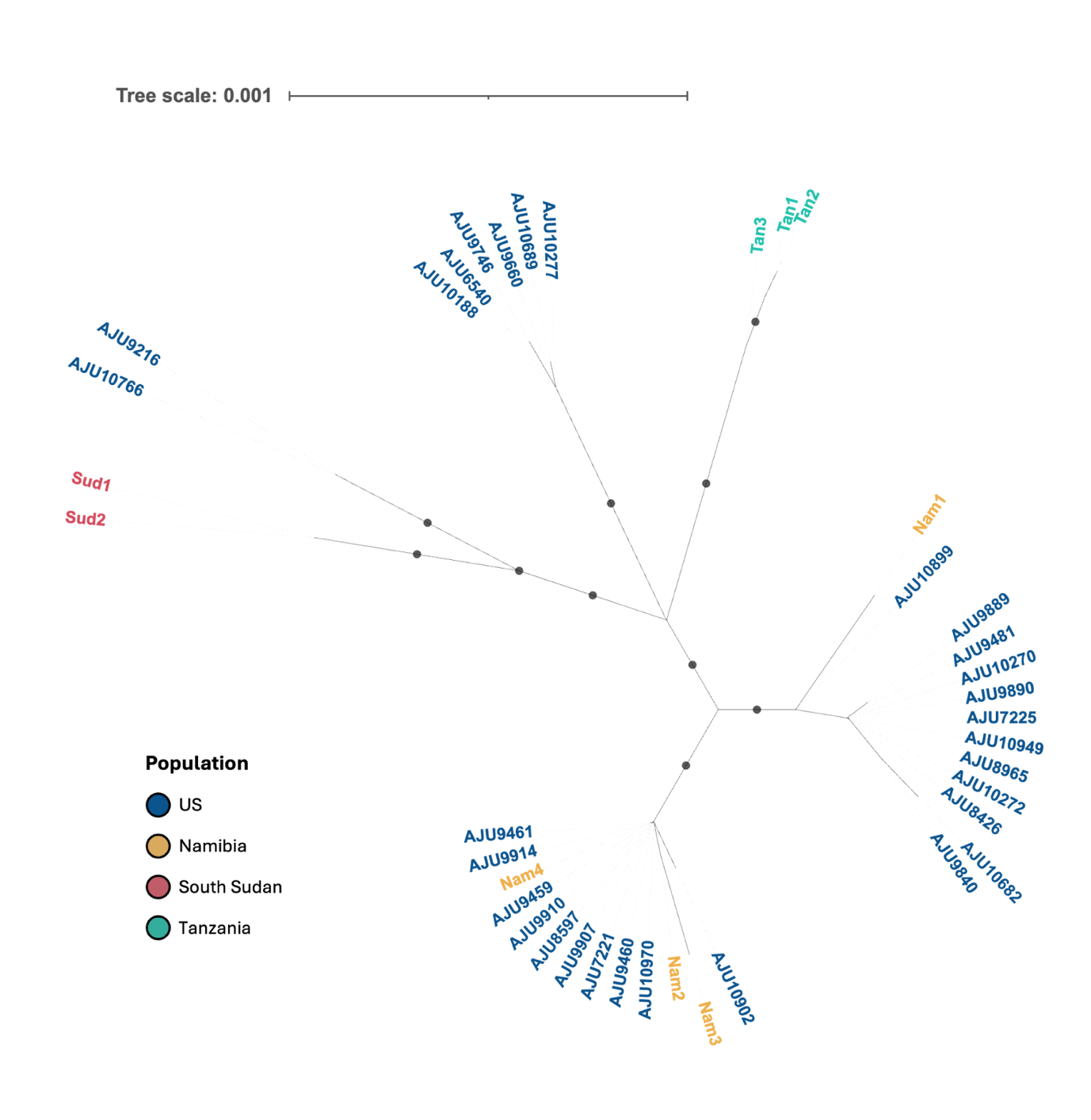

**Figure S8. Maximum likelihood of the mitogenome tree of cheetah samples.** Samples are coloured by population of origin: Namibian individuals (yellow), Tanzanian individuals (green), *ex situ* U.S. individuals (blue), and sequencing replicates (red). The scale bar indicates substitutions per site. Branches with bootstrap values > 90% are marked with a grey circle.

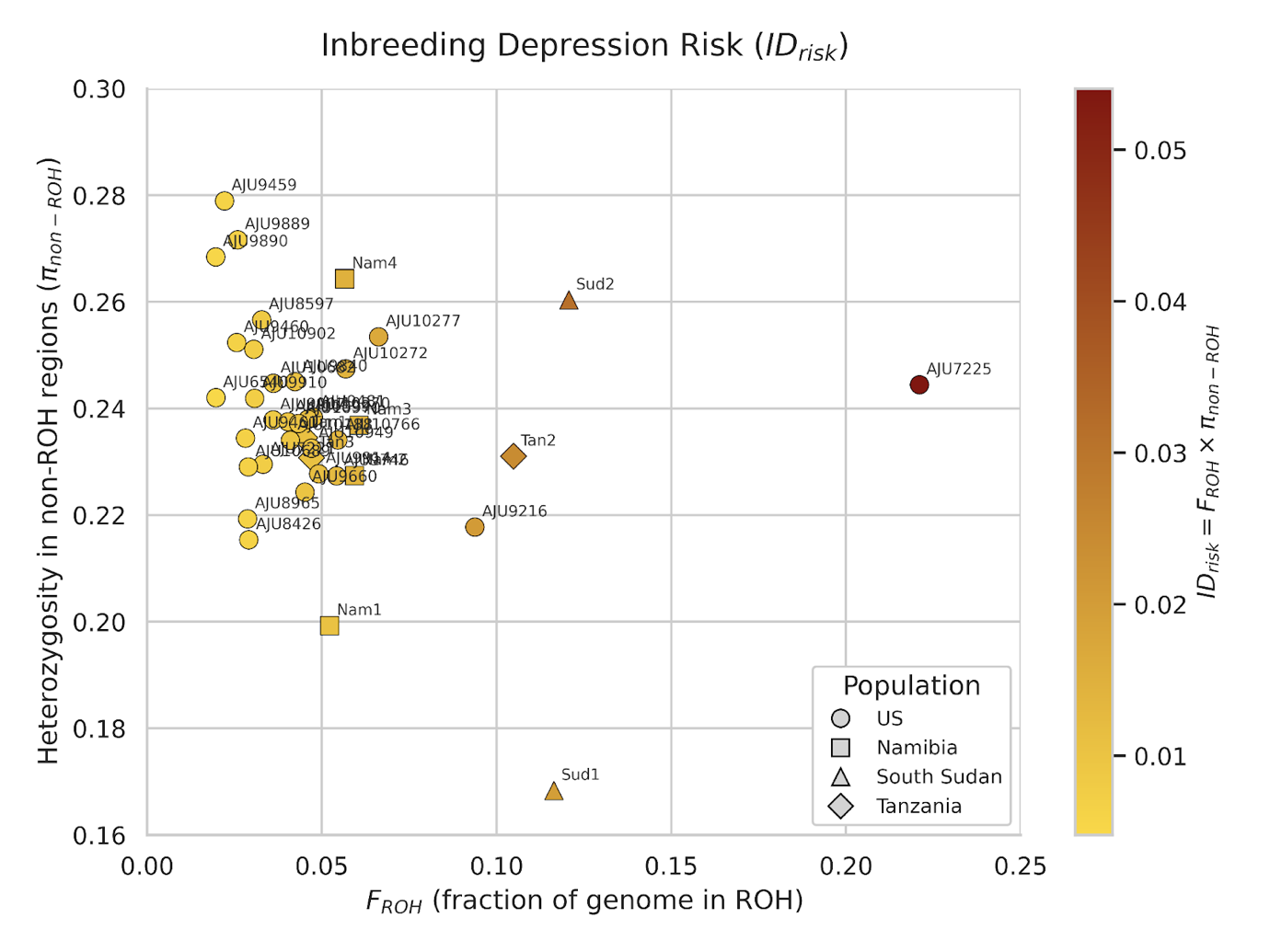

**Figure S9. Inbreeding depression risk (IDrisk) for each individual.** IDrisk scores combine a measure of the fraction of the genome in ROH (*F_ROH_*) with the average heterozygosity in non-ROH regions of the genome. Each individual is coloured based on the severity of the score, with dark red scores showing highest IDrisk. As per the thresholds published by (Kyriazis et al., 2025), all individuals show low risk of inbreeding depression. Shapes of plot points correspond to populations: U.S. (circle), Namibia (square), South Sudan (triangle) and Tanzania (diamond).

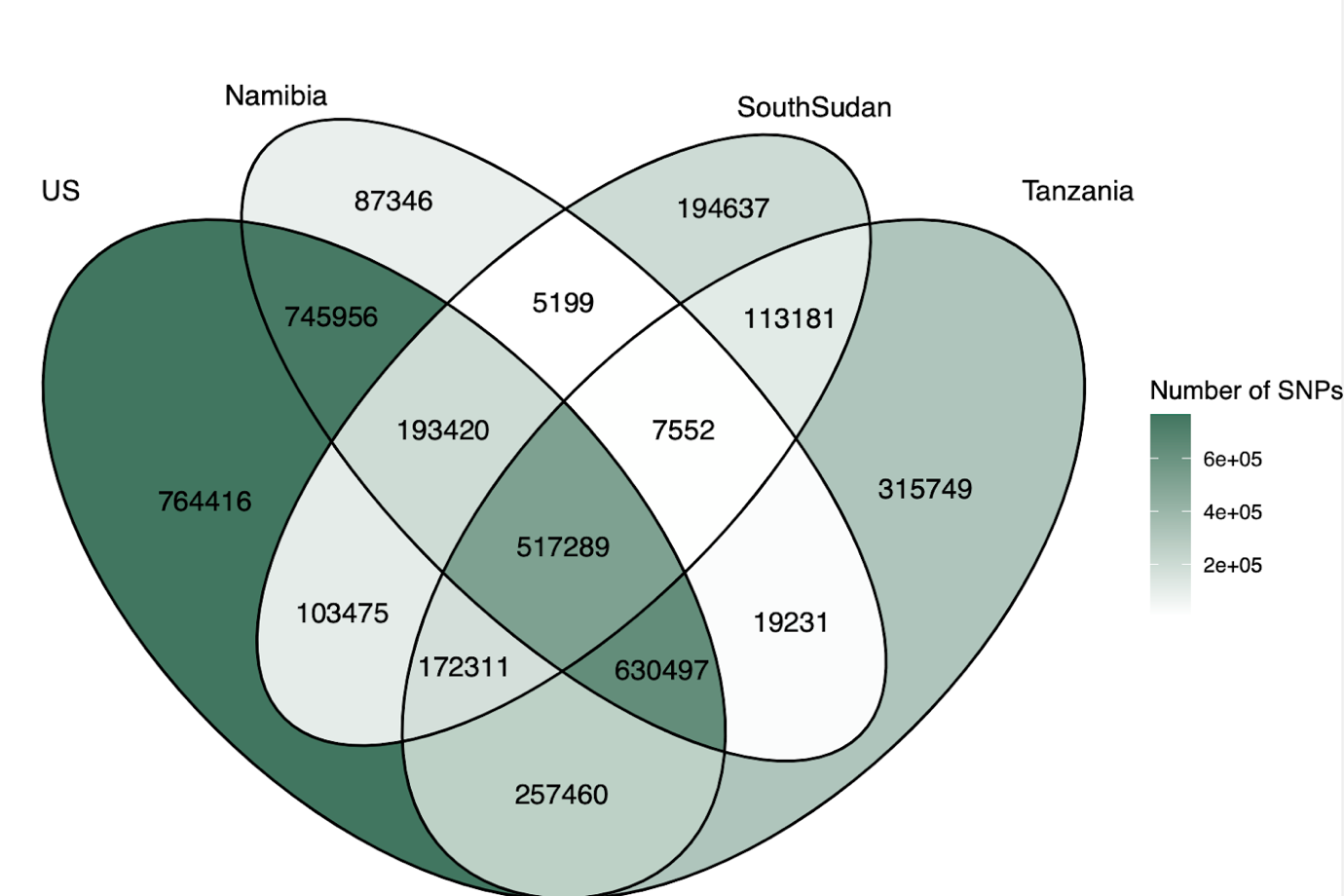

**Figure S10. Distribution of biallelic SNPs across the four cheetah populations.** The colour of each segment indicates the number of SNPs shared between those populations, with darker green indicating a higher number.

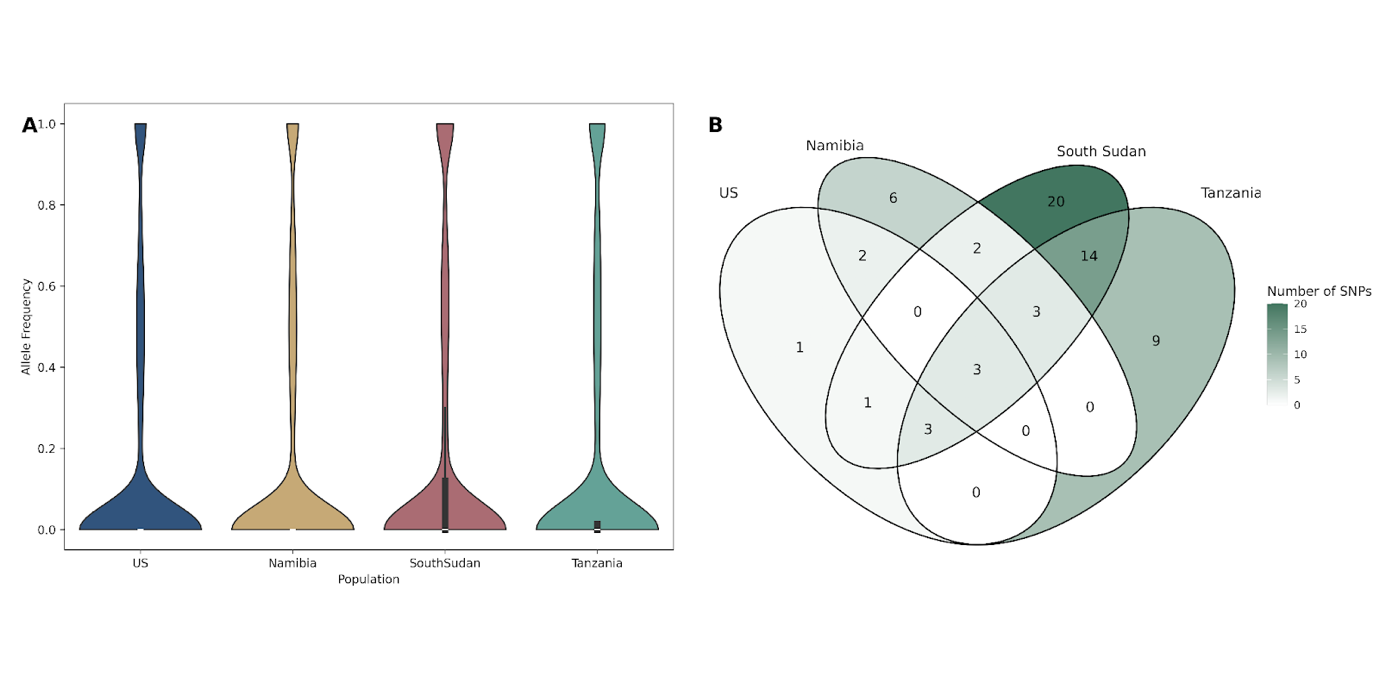

**Figure S11. Allele frequency of high-impact SNPs found in protein-coding genes.** **A** Violin plot showing the allele frequency of all high-impact SNPs per population. **B** Distribution of high-impact SNPs with an allele frequency > 0.9 across the populations. A darker colour indicates a higher number of SNPs.

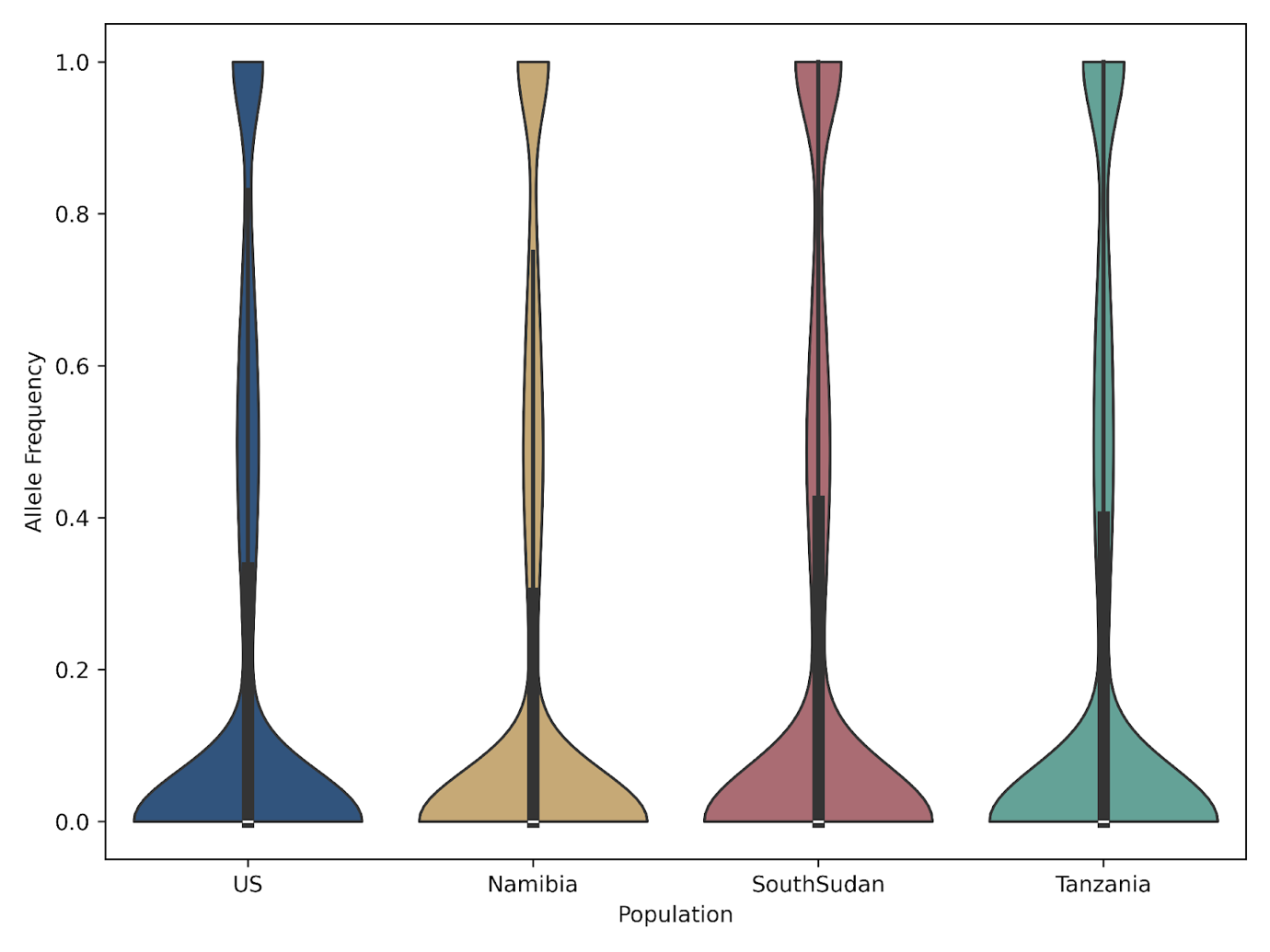

**Figure S12. Allele frequencies of moderate-impact SNPs in protein-coding genes per population.** Allele frequencies were calculated for each SNP classed as "moderate" impact by SnpEff within each population: U.S. (blue), Namibia (yellow), South Sudan (red), and Tanzania (green).

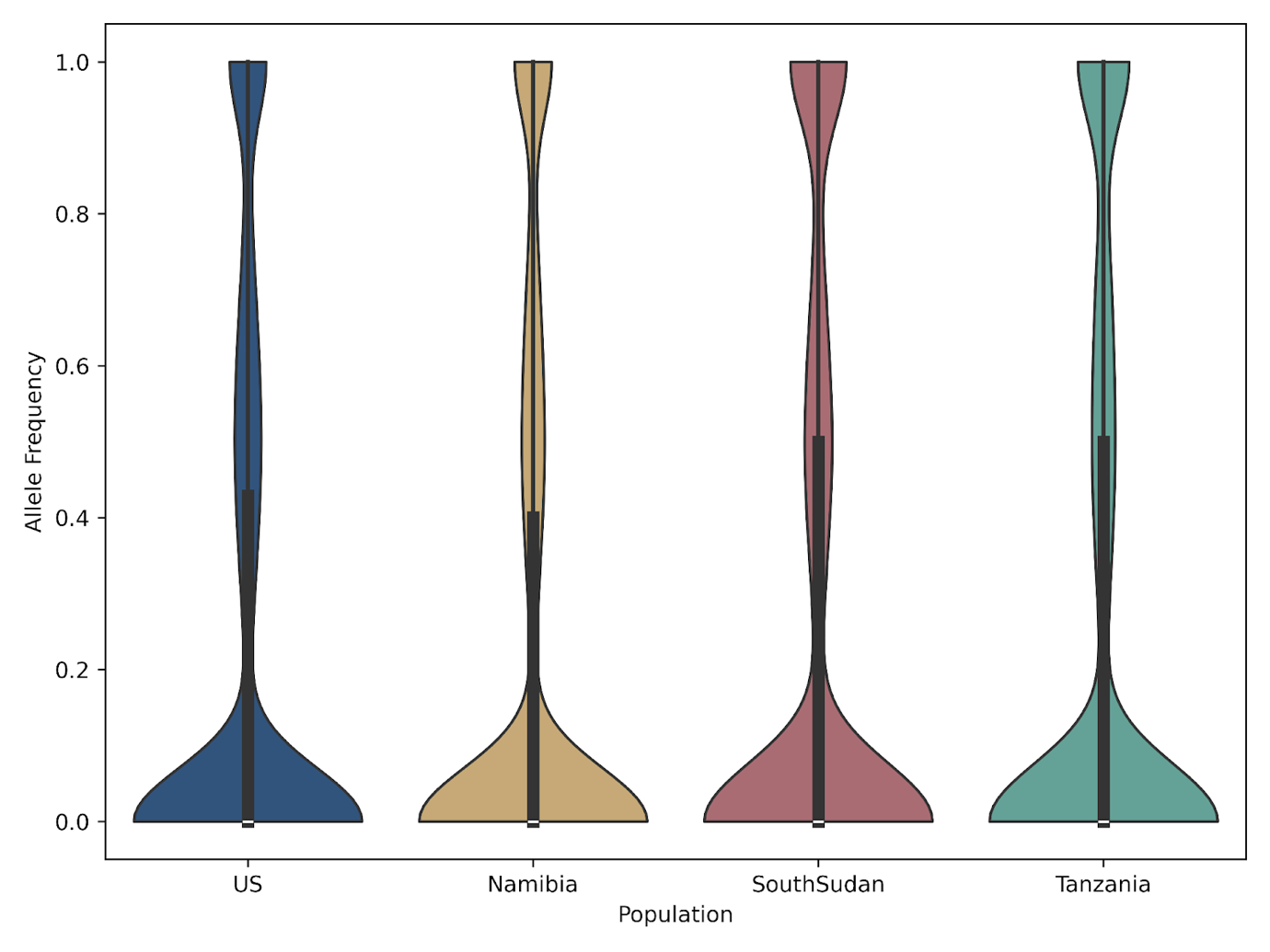

**Figure S13. Allele frequencies of low-impact SNPs in protein-coding genes per population.** Allele frequencies were calculated for each SNP classed as "low" impact by SnpEff within each population: U.S. (blue), Namibia (yellow), South Sudan (red), and Tanzania (green).

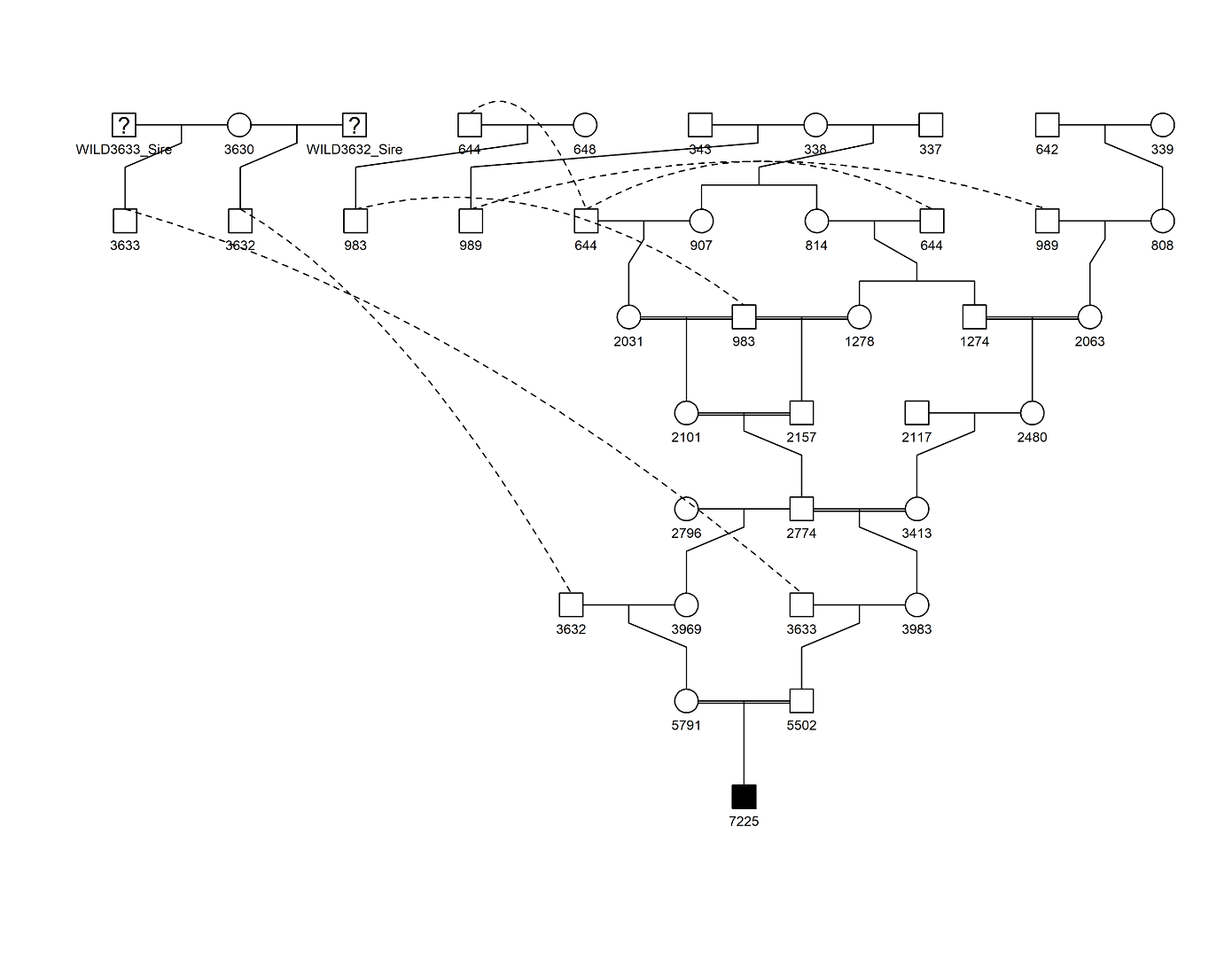

**Figure S14. Pedigree of AJU7225 from the *ex situ* U.S. population.** Males are shown with squares and females with circles. Dotted lines indicate individuals that are present in multiple locations across the pedigree. Number IDs for each individual correspond to the International Cheetah Studbook. Individuals with wild origin are denoted with "WILD"; all other individuals were born in captivity. Severe inbreeding can be observed between multiple sets of AJU7225’s ancestors.

Supplementary Table legends

**Table S1: Sample metadata.** Metadata for all cheetah samples used in this study. For all samples, their sex, source population, owner/provider is provided. For relevant samples, their house name, collection date, studbook ID and accession is also provided. *Ex situ* samples were collected by the NZCBI Veterinary Staff and provided by Dr. Adrienne Crosier and Dr. Klaus-Peter Koepfli (Smithsonian’s National Zoo and Conservation Biology Institute). South Sudan samples were collected by Kelsey Greene (African Parks South Sudan) and sequenced by Dr. Ellie Armstrong at University of California, Riverside. Owner/Providers are as follows: White Oak Conservation, Smithsonian's National Zoo and Conservation Biology Institute (NZCBI), San Diego Zoo, Fossil Rim Wildlife Centre, Wildlife Safari Winston, Toronto Zoo, Toledo Zoo.

**Table S2: Read quality and mapping.** For each sample, the total number of raw sequence reads is provided, followed by the number and percentage of base pairs removed through quality trimming (using Trim_galore) and the remaining read numbers. The number and percentage of reads that mapped successfully (using BWA-MEM) is shown alongside the average sequencing depth for each individual.

**Table S3: Significance values for genetic diversity statistics.** For each population statistic, a Kruskal-Wallis test was carried out to determine overall significance. H and p-values are provided. For each pair of populations, pairwise Wilcoxon rank-sum tests were carried out, followed by Bonferroni correction of p-values.

**Table S4: Significance values for genetic load.** For each combination of masked and realized load of high- and moderate-impact SNPs, a Kruskal-Wallis test was carried out to determine overall significance. H and p-values are provided. For each pair of populations, pairwise Wilcoxon rank-sum tests were carried out, followed by Bonferroni correction of p-values.

**Table S5: GO enrichment analysis for high-impact coding SNPs.** Gene ontology (GO) enrichment analysis was run using ShinyGO v0.85. The foreground set of genes were those with a predicted high-impact SNP which also had a corresponding 1:1 ortholog in human. The background set of genes was all 1:1 orthologs between cheetah and human. Results reported are those with an enrichment false discovery rate (FDR) corrected q-value cutoff of 0.05.

**Table S6: GO enrichment analysis for moderate-impact coding SNPs.** Gene ontology (GO) enrichment analysis output was run using ShinyGO v0.85. The foreground set of genes were those with a predicted high-impact SNP which also had a corresponding 1:1 ortholog in human. The background set of genes was all 1:1 orthologs between cheetah and human. Results reported are those with an enrichment false discovery rate (FDR) corrected q-value cutoff of 0.05.

**Table S7: Population distribution of previously-identified PTCs.** 65 previously identified premature termination codon (PTC) loci (Peers et al., 2025) were extracted from the full population VCF using the  aciJub1 assembly as the reference genome (Dobrynin et al., 2015). For each locus, the position of the PTC is given, alongside the gene name, its status in the VCF, the reference and alternate alleles, site quality and the number of individuals with each genotype. The final outcome of each locus is given: each mutation is either unique to aciJub1 (the reference), variant in the population or fixed in all individuals sampled. Two sites had insufficient coverage to call.
